# WHaloForce enables chemigenetic imaging of molecular tension in living cells and animals

**DOI:** 10.64898/2026.08.18.745627

**Authors:** Di Wu, E. Angelo Morales, Joon Lee, Sushil Pangeni, Helen Farrants, Katherine A. Hutchings, Robert J. Barndt, Qi Yu, Xuesong Li, Hari Shroff, Haodi Wu, Alison G. Tebo, Luke D. Lavis, Eric R. Schreiter, Taekjip Ha, Shaohe Wang

**Affiliations:** Janelia Research Campus, Howard Hughes Medical Institute; Program in Cellular and Molecular Medicine, Boston Children’s Hospital; Department of Biological Chemistry and Molecular Pharmacology, Harvard Medical School; Department of Biophysics, Johns Hopkins University; Department of Cell Biology, Johns Hopkins University School of Medicine; Heart, Lung, and Blood Vascular Medicine Institute, University of Pittsburgh School of Medicine; Division of Cardiology, Department of Medicine, University of Pittsburgh School of Medicine; School of Life Sciences, Southern University of Science and Technology, Shenzhen, China; Howard Hughes Medical Institute

## Abstract

Piconewton forces borne by individual proteins within complex assemblies underlie cell adhesion, migration, and tissue morphogenesis, yet remain difficult to image in living systems. Here, we introduce WHaloForce, a chemigenetic HaloTag-based tension sensor that converts force-dependent relief of tryptophan-mediated dye quenching into a fluorescence lifetime change. Optical tweezers revealed a switch-like unquenching transition near 5 pN, and cellular measurements demonstrated dynamic responses to actomyosin activity. WHaloForce enabled quantitative tension imaging of diverse force-bearing proteins (vinculin, E-cadherin, α-catenin, and laminin) in mammalian cells, mouse tissue, and *C. elegans*. Bright synthetic dyes made tension measurements possible at endogenous expression levels. In *C. elegans*, vinculin and laminin showed opposite tension patterns between tissues, highlighting tissue-specific force transmission through adhesions and the extracellular matrix. We also identified ovulation as a source of molecular loading within the spermathecal laminin network. WHaloForce thus offers a modular platform for imaging spatiotemporal tension patterns in living systems.

## Introduction

Mechanical forces shape biological structures and regulate cell behavior. At the molecular scale, piconewton-range tensions borne by cytoskeletal, adhesion, and extracellular matrix (ECM) proteins underlie cell adhesion, migration, morphogenesis, muscle contraction, and mechanotransduction^1–4^. Yet these forces are difficult to measure in their native context: they act on specific proteins within crowded assemblies, change in magnitude and direction over time, and are often buried deep inside living tissue. Reading molecular tension in situ therefore requires genetically encoded reporters that convert force into an optical signal without disrupting the mechanical linkages they are designed to study.

Genetically encoded molecular tension sensors have transformed the study of mechanobiology, but existing approaches have important tradeoffs. FRET-based sensors place an extensible linker between donor and acceptor fluorophores, so that force-induced extension reduces energy transfer^5^. Since their introduction in vinculin^6,7^, these probes have enabled calibrated and reversible measurements of tension across many proteins^8–13^. However, FRET sensors require two fluorescent proteins, increasing probe size and occupying two spectral channels; the resulting FRET changes are often modest. A complementary strategy uses force-dependent exposure of cryptic motifs to recruit fluorescent probes, converting molecular tension into a high-contrast localization or intensity signal^14,15^. These sensors can reveal mechanically induced conformational states, but their readout depends on local probe concentration, recruitment kinetics, and binding equilibria; in some cases, recruitment may also stabilize the exposed state^15^. Together, these architectural and readout constraints limit quantitative, protein-specific tension imaging at endogenous expression levels in intact tissues and living animals, particularly when additional fluorescent labels are needed.

Recent work on chemigenetic indicators suggests an alternative strategy. WHaloCaMP is a HaloTag-based calcium indicator in which a calcium-sensing module is inserted into a rhodamine dye-labeled HaloTag scaffold^16,17^. In the calcium-free state, a strategically positioned tryptophan quenches the bound rhodamine dye through photoinduced electron transfer (PET). Calcium binding alters the sensor geometry, relieving quenching and increasing both fluorescence intensity and lifetime^16^. This design highlights the power of coupling a ligand-responsive protein module to a compact, covalently labeled dye scaffold, thereby converting a conformational change into a switch in the dye’s emissive state. Because the signal arises from one bright synthetic fluorophore rather than a donor-acceptor pair or recruited binding partner, tryptophan-rhodamine PET quenching provides an attractive but previously unexplored opportunity for chemigenetic molecular tension sensing.

Here, we introduce WHaloForce, a chemigenetic molecular tension sensor that repurposes tryptophan-rhodamine PET quenching from chemical sensing to mechanical sensing. Built from a circularly permuted HaloTag engineered for force-dependent unquenching of a bound rhodamine dye, WHaloForce converts piconewton-scale mechanical load into a change in the dye’s fluorescence lifetime. Because lifetime is largely independent of probe concentration, excitation power, and labeling density, fluorescence lifetime imaging microscopy (FLIM) remains quantitative even when these experimental conditions vary^18,19^. Single-molecule measurements showed a switch-like transition near 5 pN, well matched to the low-piconewton forces commonly borne by individual proteins in cellular force-transmission pathways^6–12^. Inserted into vinculin, cadherincatenin junctional proteins, and laminin, WHaloForce enabled molecular tension imaging in cultured cells and, through endogenous CRISPR knock-ins, in mouse tissue and living *C. elegans*. Across these systems, WHaloForce resolved tension patterns that varied in space, in time, and between proteins within the same tissue. Together, these results establish WHaloForce as a compact chemigenetic platform for FLIM-based molecular tension imaging in cells, tissues, and living animals.

## Results

### cpWHalo undergoes a switch-like, force-dependent unquenching transition near 5 pN

To test whether the tryptophan-mediated PET quenching strategy used in WHaloCaMP could be adapted for force sensing, we engineered a circularly permuted HaloTag (cpHalo) by connecting the original C-terminus to the original N-terminus with a flexible linker and introducing new termini at residue R179, the insertion site of the calcium-sensing module in WHaloCaMP^16^, along with a P180V substitution at the new N-terminus (**Fig. 1a**). To enable quenching, cpWHalo carries G171W, which places a tryptophan adjacent to the bound dye, together with V178W (**Fig. 1a,b**; **Supplementary Note 1**). P180V, G171W, and V178W were all carried over from earlier HaloTag engineering.

**Figure 1.**
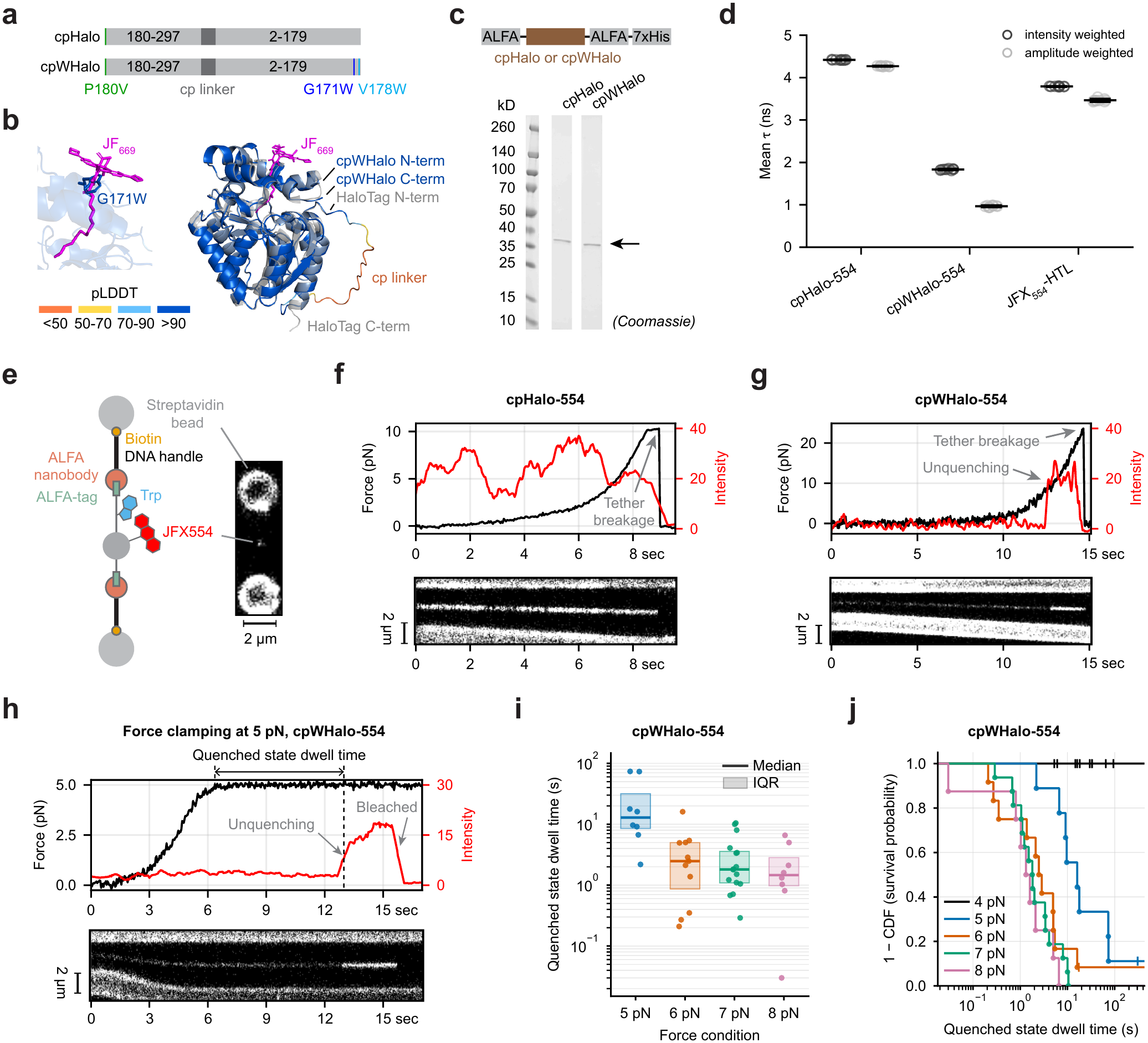
cpWHalo undergoes a force-dependent unquenching transition. **(a)** Schematics of cpHalo and cpWHalo design. **(b)** AlphaFold-predicted structure of cpWHalo, colored by pLDDT score, overlaid with the crystal structure of HaloTag7-JF_669_ (PDB: 8SW8). The left zoom-in highlights the proximity of the engineered quenching tryptophan, G171W, in cpWHalo to the rhodamine dye position in HaloTag7-JF_669_. **(c)** Schematic of ALFA-tagged cpHalo and cpWHalo proteins used for single-molecule tethering (top) and Coomassie-stained SDS-PAGE gel of the purified proteins (bottom). **(d)** Intensity-weighted and amplitude-weighted mean fluorescence lifetimes (τ) for the indicated protein-dye conjugates and the free dye-HaloTag ligand (HTL) control. **(e)** Schematic of the single-molecule tethering geometry (left) and representative image of a cpWHalo-554 tether between two beads (right). **(f–h)** Representative smoothed force and fluorescence intensity traces (top) and corresponding kymographs (bottom) for cpHalo-554 or cpWHalo-554. cpHalo-554 was measured during force ramping **(f)**, cpWHalo-554 during force ramping **(g)**, and cpWHalo-554 during force clamping **(h).** **(i)** Box-and-scatter plot of quenched-state dwell times for cpWHalo-554 under the indicated constant-force conditions. The y-axis is plotted on a log scale. IQR: interquartile range. The 4 pN condition is not shown because no unquenching transitions were observed. **(j)** Survival probability plot of cpWHalo-554 quenched-state dwell times under the indicated force conditions. Dots indicate observed unquenching transitions, and vertical tick marks indicate right-censored traces that ended because of photobleaching, tether breakage, or the planned end of the recording. The x-axis is plotted on a log scale.

In HEK293T cells, cpHalo-dye conjugates showed fluorescence lifetimes comparable to HaloTag but reduced brightness across all 17 dyes tested^20–23^ (**Extended Data Fig. 1**). These two readouts report different photophysics. Rhodamine dyes interconvert between a colorless lactone and a fluorescent zwitterion, and shifting this equilibrium changes absorbance, and therefore brightness, while leaving lifetime unchanged^20–23^. cpHalo therefore appears to shift the equilibrium of the bound dye toward the non-absorbing lactone. By contrast, cpWHalo-dye conjugates showed reduced lifetime as well as brightness relative to cpHalo (**Extended Data Fig. 1**), a signature of tryptophan-mediated PET quenching, which opens a nonradiative decay pathway from the excited state and lowers quantum yield and lifetime together^19^. We selected JFX_554_ for further analysis because it combined a large lifetime dynamic range between cpHalo and cpWHalo (Δτ_int_ = 3.0 ns; 3.3-fold) with a modest 2.5-fold brightness reduction (**Extended Data Fig. 1**).

To evaluate force responsiveness, we purified cpHalo and cpWHalo with ALFA tags appended to both termini to enable downstream tethering in single-molecule experiments (**Fig. 1c**), and we labeled both proteins with JFX_554_-HaloTag ligand (HTL). We first confirmed in bulk that the engineered quenching was retained in the purified, soluble proteins: cpWHalo-554 showed a 4.4-fold reduction in amplitude-weighted fluorescence lifetime relative to cpHalo-554 (**Fig. 1d**). This lifetime reduction corresponded to a 4.5-fold decrease in quantum yield with only an 8.5% change in extinction coefficient (**Extended Data Fig. 2a,b**), confirming that tryptophan-mediated PET quenching primarily affects quantum yield rather than absorption, as expected for a PET mechanism. The close agreement between the decreases in quantum yield and fluorescence lifetime is consistent with predominantly dynamic quenching under these solution conditions^24^.

We then tethered single molecules of cpWHalo-554 or cpHalo-554 between two streptavidin-coated beads via dsDNA handles functionalized with biotin and ALFA-tag nanobodies^25^, enabling simultaneous application of calibrated force and measurement of fluorescence (**Fig. 1e**). Upon gradual increases in applied force, cpWHalo-554, but not cpHalo-554, exhibited abrupt fluorescence increases (**Fig. 1f,g**), consistent with a discrete transition from a quenched to an unquenched state. This behavior likely reflects a force-induced structural rearrangement that disrupts the geometric coupling required for tryptophan-mediated PET quenching.

We quantified the force dependence of this transition by measuring the unquenching rate (k_QU_) under constant force. At 4 pN, no unquenching events were observed within the experimental time window, placing an upper bound of k_QU_ < 0.0106 s^-1^. At 5 pN, unquenching occurred stochastically on a ∼64-s timescale (k_QU_ = 0.0156 s^-1^). At 6–8 pN, transitions often occurred within 6 s (k_QU,6pN_ = 0.188 s^-1^, k_QU,7pN_ = 0.311 s^-1^, k_QU,8pN_ = 0.433 s^-1^). Thus, although individual transitions remained stochastic, the unquenching rate increased sharply above ∼5 pN, corresponding to a 12-fold increase between 5 and 6 pN (**Fig. 1h–j**). This steep force-dependent acceleration indicates that WHaloForce response kinetics are load-dependent, providing a plausible basis for rapid lifetime changes when molecular tension transiently exceeds the ∼5-pN transition range in cells.

To test reversibility, we reduced the force after an unquenching transition. In two tethers, fluorescence returned to the low state (**Extended Data Fig. 2c,d**). In one of these, reapplying 7 pN left the molecule in the low-intensity state for several seconds before it underwent a second unquenching transition (**Extended Data Fig. 2c**). Because the tether was again under high tension, the persistence of the low-intensity state indicates that it corresponds to a bona fide quenched state rather than a detection or geometric artifact. Comprehensive characterization of re-quenching was limited by photobleaching at the high excitation intensities required for single-molecule detection. These traces establish that force-induced unquenching can reverse, but do not determine the frequency, completeness, or kinetics of re-quenching across the sensor population.

Together, these results establish cpWHalo as a force-responsive HaloTag variant with a switch-like unquenching transition near 5 pN, within the range of molecular tensions commonly experienced by proteins in cells^6–12,26–29^, supporting its use as a tension sensor. Because the sensor occupies one of two quenching states, the mean lifetime across a cell or region shifts with the proportion of sensors in the unquenched state rather than reporting a single force value. Its temporal relationship to force therefore depends on both unquenching and re-quenching kinetics. Hereafter, we refer to cpWHalo as WHaloForce in tension-sensing constructs, including terminal fusion controls.

### WHaloForce reports heterogeneous and dynamic vinculin tension at focal adhesions

To evaluate WHaloForce in cells, we inserted it into vinculin, a focal adhesion protein that transmits forces between integrins and the F-actin cytoskeleton^2,30^. WHaloForce was inserted between the vinculin head and tail domains after residue E883, the same position used in a previous FRET-based tension sensor^6^, generating Vin-WF. As controls, we generated Vin-cpH, which contains cpHalo without the quenching tryptophan, and Vin-N-WF, in which WHalo-Force was fused to the N-terminus outside the primary force-transduction axis (**Fig. 2a**). Throughout this study, [protein]-WF denotes an internal WHaloForce insertion, [protein]-cpH denotes the corresponding non-quenching cpHalo control, and [protein]-N/C-WF denotes a terminal, off-axis WHaloForce fusion.

**Figure 2.**
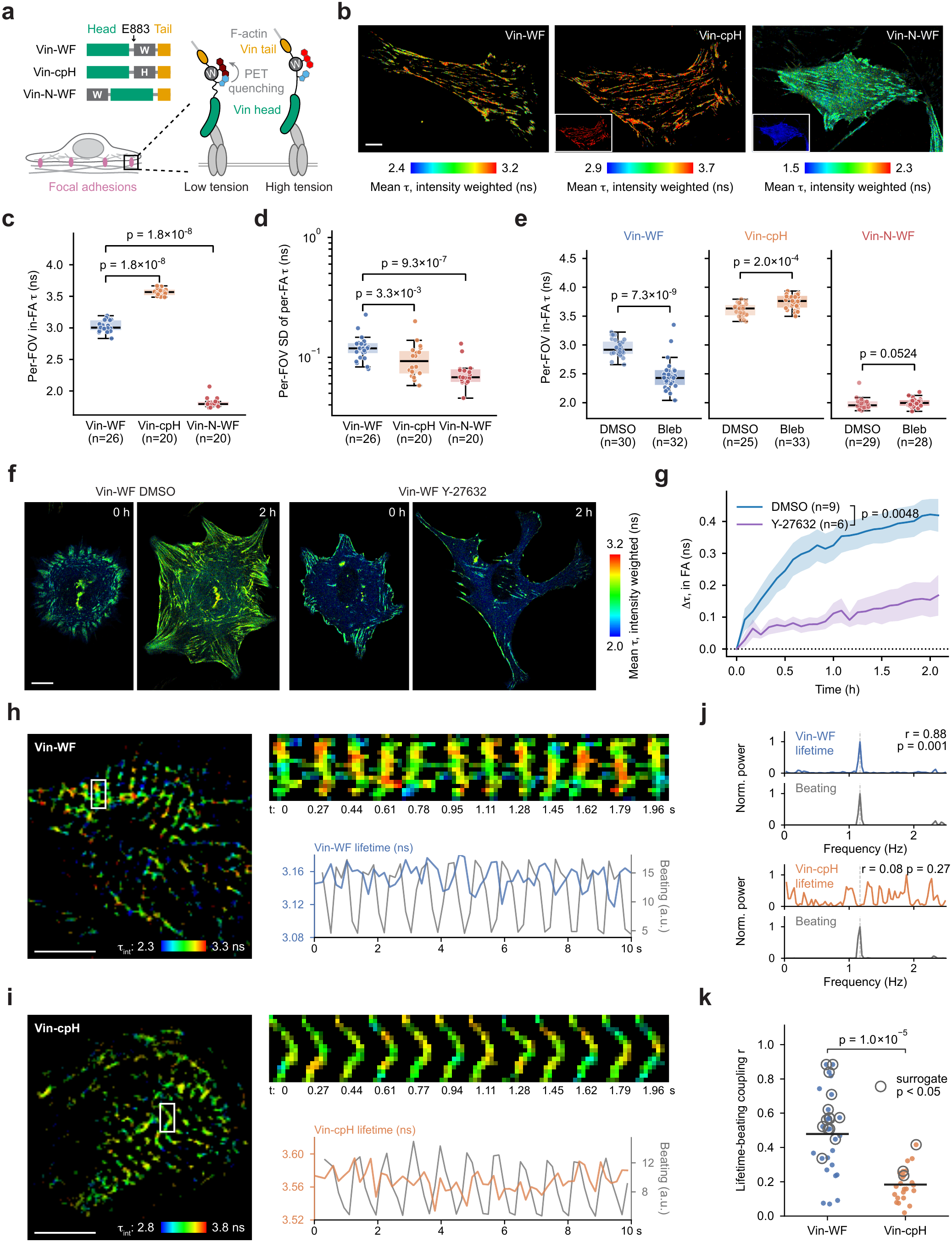
WHaloForce reports heterogeneous and dynamic vinculin tension at focal adhesions. **(a)** Schematics of Vin-WF, Vin-cpH, and Vin-N-WF sensor and control designs, and model of low- and high-tension Vin-WF states at focal adhesions (FAs). **(b)** FLIM images of NIH-3T3 cells expressing Vin-WF, Vin-cpH, or Vin-N-WF. Color represents intensity-weighted mean fluorescence lifetime (τ). Color scales were adjusted independently for each image to avoid saturation. Insets show Vin-cpH and Vin-N-WF displayed using the same color scale as Vin-WF, resulting in saturation, to facilitate comparison across constructs. **(c–e)** Box-and-scatter plots of per-field-of-view (FOV) mean in-FA intensity-weighted lifetime **(c,e)** or per-FOV standard deviation (SD) of per-FA intensity-weighted lifetime **(d)**; each point represents one FOV, and different FOVs contain different cells. Cells in **(c,d)** were untreated. Cells in **(e)** were treated with 20 μM blebbistatin or an equal volume of DMSO for 12 h. Statistics: Mann-Whitney U tests with Holm-Bonferroni correction; reported p values are Holm-adjusted. **(f)** FLIM images of NIH-3T3 cells expressing Vin-WF during cell spreading under DMSO control conditions or after treatment with the ROCK inhibitor Y-27632 (5 μM), shown at 0 and 2 h as indicated. **(g)** Change in Vin-WF fluorescence lifetime relative to the starting time point for DMSO control and 5 μM Y-27632-treated NIH-3T3 cells during spreading. Shading represents SEM. p value was determined by Mann-Whitney U test comparing area under the curve (AUC) between conditions. n, cells. **(h,i)** FLIM image at the starting time point (left), time series (top right), and corresponding traces of mean intensity-weighted lifetime in bright costameres and beating motion (bottom right) for human iPSC-derived cardiomyocytes expressing Vin-WF **(h)** or Vin-cpH **(i)**. Beating motion was quantified as the mean absolute frame-to-frame intensity difference within the focal-adhesion region.**(j)** Normalized power spectra (Fourier analysis) of beating traces (gray) and bright-costamere lifetime traces (colored) for representative cells of Vin-WF (top) and Vin-cpH (bottom). Dashed line indicates the beat fundamental frequency, f_0_. r indicates lifetime-beating coupling calculated by lag-optimized cross-correlation within ±1/2 beat cycle; p indicates significance by phase-randomized surrogate testing (1,000 randomizations).**(k)** Lifetime-beating coupling r, calculated by lag-optimized cross-correlation within ±1/2 beat cycle, for individual cells of Vin-WF (n=27) and Vin-cpH (n=23). Open circles indicate cells with significant coupling by phase-randomized surrogate testing (1,000 randomizations, p < 0.05). Bars indicate construct means. The p value shown compares r values between constructs (two-sided Mann-Whitney U test). The fraction of cells with significant coupling (14 of 27 versus 3 of 23) was compared separately by two-sided Fisher’s exact test (p = 0.006). All cells were labeled with JFX_554_-HTL. Scale bars, 10 μm.

In NIH-3T3 fibroblasts, Vin-WF and the control constructs localized to focal adhesions (**Fig. 2b**). Vin-WF exhibited substantial fluorescence lifetime heterogeneity between focal adhesions within the same cell (**Fig. 2c,d**), suggesting heterogeneous tension loading across vinculin molecules. In contrast, Vin-cpH showed uniformly high lifetimes, as expected for a non-quenched cpHalo module, whereas the off-axis control Vin-N-WF showed low, spatially uniform lifetimes across focal adhesions (**Fig. 2b–d**). Importantly, inhibiting myosin contractility reduced focal adhesion size and density in all constructs but shortened fluorescence lifetimes only in Vin-WF; Vin-N-WF was unchanged, and Vin-cpH showed a small increase (**Fig. 2e, Extended Data Fig. 3a–c**). Together, these results indicate that Vin-WF reports spatially heterogeneous, actomyosin-dependent tension borne by vinculin at focal adhesions.

We next performed live imaging to examine vinculin tension dynamics in spreading NIH-3T3 cells under control conditions and after ROCK inhibitor treatment. In control cells, Vin-WF fluorescence lifetimes increased as focal adhesions matured, whereas this increase was significantly blunted by ROCK inhibition, further indicating that force transmission through vinculin depends on myosin contractility (**Fig. 2f,g; Video S1**).

We next examined Vin-WF dynamics in human iPSC-derived cardiomyocytes (iCMs), which undergo spontaneous contractions. iCMs contain specialized focal adhesions called costameres that mechanically couple sarcomeres to the substrate^31,32^ (**Fig. 2h**). During spontaneous beating, costameres exhibited rhythmic displacements, and Vin-WF lifetimes fluctuated in step with them (**Fig. 2h**; **Video S2**). Fourier analysis identified a frequency component in the lifetime traces that matched the beating rhythm (**Fig. 2j**), and the lifetime-beating coupling was significant in 14 of 27 cells (**Fig. 2k**). The force-insensitive control Vin-cpH, which lacks the quenching tryptophan, showed no such structure: its lifetime fluctuations carried no frequency component at the beating rhythm (**Fig. 2i,j**; **Video S3**) and were coupled in only 3 of 23 cells, consistent with detection noise expected when FLIM is pushed to the acquisition speeds needed to resolve the beating rhythm (**Fig. 2k**; Fisher’s exact test vs Vin-WF, p = 0.006). These data indicate that Vin-WF lifetimes at costameres can track the beating rhythm of iCMs (∼1.2 Hz), supporting a dynamic component of the sensor response in this context.

Collectively, these results show that WHaloForce reports heterogeneous and dynamic vinculin tension at focal adhesions. In fibroblasts and beating cardiomyocytes, Vin-WF resolved spatial tension differences and second-scale life-time fluctuations.

### WHaloForce reveals anisotropic and asymmetric tension at epithelial junctions

To test whether the WHaloForce strategy could be generalized to other force-transducing proteins, we inserted the sensor into E-cadherin and α-catenin, two core components of adherens junctions in epithelial cells. E-cadherin is a transmembrane adhesion protein whose extracellular domain mediates homophilic interactions between neighboring cells, while its intracellular domain connects to F-actin through β-catenin and α-catenin^33^. For E-cadherin, we inserted WHaloForce or cpHalo after residue A780, between the juxtamembrane domain (JMD) and the catenin-binding domain (CBD), generating Ecad-WF and the Ecad-cpH control (**Fig. 3a; Extended Data Fig. 4a,b**). For α-catenin, we inserted WHaloForce or cpHalo after residue T654, within the linker between the M domain and the C-terminal F-actin-binding domain (FABD)^34–36^, yielding aCat-WF and aCat-cpH (**Fig. 3a; Extended Data Fig. 4c–e**). For both proteins, we also generated C-terminal WHaloForce fusions, Ecad-C-WF and aCat-C-WF, to place the sensor outside the primary force-transducing axis (**Fig. 3a**).

**Figure 3.**
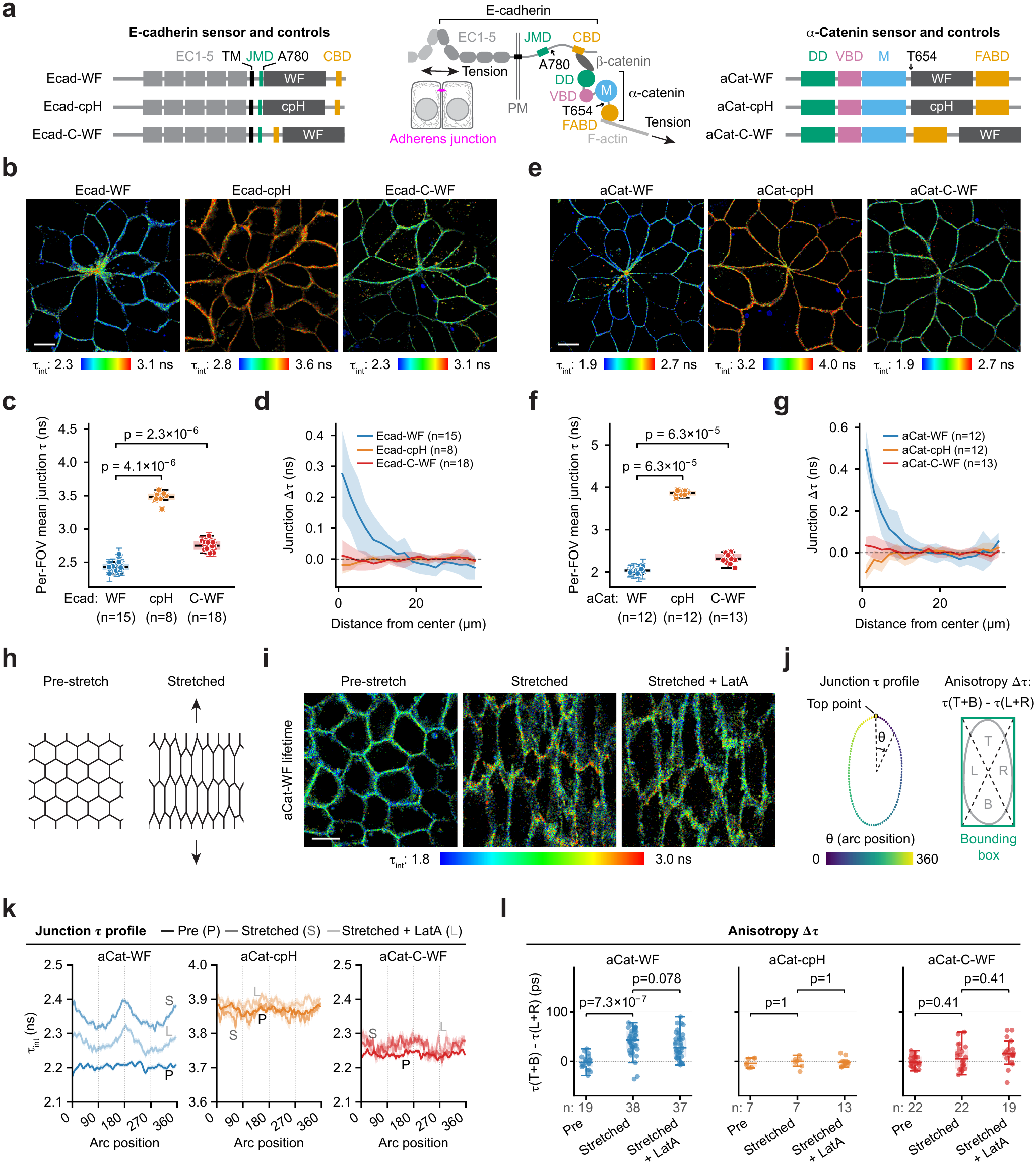
WHaloForce reveals anisotropic and asymmetric tension at epithelial junctions. **(a)** Schematics of sensor and control designs for E-cadherin (left) and α-catenin (right), with a model of the force-transmission axis at adherens junctions (middle). **(b,e)** FLIM images of MDCKII cells expressing E-cadherin sensors and controls **(b)** or α-catenin sensors and controls **(e)**. Color represents intensity-weighted mean fluorescence lifetime (τ_int_). Color scales were adjusted independently for each image to avoid saturation. **(c,f)** Box-and-scatter plots of per-field-of-view (FOV) mean junction lifetime for E-cadherin **(c)** and α-catenin **(f)** sensors and controls. Each point represents one FOV; error bars around each point indicate the within-FOV SD across junction segments included in the analysis (≥2 μm, ≥5,000 photons). Boxes show the distributions of per-FOV mean lifetimes. Statistics: Mann-Whitney U tests with Holm-Bonferroni correction; reported p values are Holm-adjusted. **(d,g)** Radial profiles of junction lifetime change (Δτ) from the rosette center for E-cadherin **(d)** and α-catenin **(g)** constructs. Δτ was calculated as the per-ring mean junction lifetime minus the baseline lifetime for the same FOV, defined as the mean junction lifetime at r ≥ 10 μm. Lines indicate means across FOVs; shaded bands indicate ±SD. n indicates the number of FOVs per construct. **(h)**Schematic of junction orientation before and after uniaxial stretch. **(i)**FLIM images of MDCKII cells before stretch (left), after stretch (middle), and after 1 μM latrunculin A (LatA) treatment (right). Color represents intensity-weighted mean fluorescence lifetime (τ_int_). **(j)**Schematics of the junction lifetime profile analysis (left) and anisotropy analysis (right). Junction lifetime profiles were generated by tracing adherens junctions around the cell perimeter, starting from the top point of each cell and proceeding clockwise. Anisotropy Δτ was calculated as the difference between the mean lifetime of pooled junction pixels in the top and bottom regions and the mean lifetime of pooled junction pixels in the left and right regions; regions were defined by the diagonal lines of each cell-bounding box. **(k)**Junction lifetime profiles for the indicated constructs and conditions, analyzed as defined in **(j)**. Lines indicate means across FOVs; shaded bands indicate ±SEM. **(l)**Junctional lifetime anisotropy Δτ for the indicated constructs and conditions, analyzed as defined in (j). Each point represents one FOV. Boxes show the interquartile range with median lines; whiskers extend to the most extreme values within 1.5 times the interquartile range. Statistics: two-sided Mann-Whitney U tests with Holm-Bonferroni correction across the two displayed comparisons within each construct; reported p values are Holm-adjusted. All cells were labeled with JFX_554_-HTL. Scale bars, 10 μm.

When expressed in MDCKII epithelial cells, Ecad-WF, aCat-WF, and their control constructs localized to cell-cell junctions as expected (**Fig. 3b,e**). At confluence, dying MDCKII cells undergo apical extrusion^37^, leaving behind rosette-like structures formed by neighboring cells. Near the rosette center, both Ecad-WF and aCat-WF displayed locally elevated fluorescence lifetimes that were absent in their corresponding cpHalo or C-terminal WHaloForce controls (**Fig. 3b–g; Extended Data Fig. 4f–i**), consistent with higher tension across adhesion molecules at these multicellular junctions^38^. Notably, Ecad-C-WF and aCat-C-WF showed higher basal lifetimes than Ecad-WF and aCat-WF, but their signals were spatially uniform at cell-cell junctions (**Fig. 3b–g**). Thus, the elevated lifetimes of the C-terminal controls likely reflect reduced intrinsic quenching efficiency in this fusion geometry rather than force-dependent unquenching.

To examine how junctional tension responds to larger external mechanical loads, we grew MDCKII cells on a thin PDMS film mounted on a custom-built uniaxial stretcher^39^ (**Extended Data Fig. 5a**). The film was stretched to twice its resting length along one axis (100% engineering strain), with concomitant contraction along the perpendicular axis. Upon stretch, aCat-WF, but not the control constructs, showed increased fluorescence lifetimes specifically at junctions oriented perpendicular to the stretch direction (**Fig. 3h–l; Extended Data Fig. 5b–d**). This indicates that α-catenin experiences anisotropic tension increases as epithelial junctions resist external deformation. The lifetimes of aCat-WF, but not the controls, decreased upon acute treatment with latrunculin A, an actin polymerization inhibitor (**Fig. 3i–l**). However, latrunculin A treatment after stretching did not reduce the lifetimes to the basal level prior to stretching, and only marginally decreased the anisotropy within the experimental time frame (30-60 min after treatment) (**Fig. 3k,l**), suggesting that mechanically loaded F-actin networks may be more resistant to acute depolymerization.

Together, these results demonstrate that WHaloForce can be adapted to multiple components of the cadherin-catenin complex, revealing asymmetric junctional tension near rosette centers following apical extrusion and anisotropic tension increases under external stretch.

### Endogenous WHaloForce knock-ins reveal tissue- and orientation-dependent tension patterns in *C. elegans*

Measuring tension in intact animals places the greatest demand on a concentration-independent readout, because expression level, labeling efficiency, imaging depth, and local optical properties all vary across tissues. To enable endogenous tension measurements in living animals, we first used CRISPR to generate knock-in *C. elegans* strains by inserting WHaloForce into endogenous loci encoding the worm orthologs of vinculin (*deb-1*), E-cadherin (*hmr-1*), and laminin γ1 (*lam-2*). These proteins transmit mechanical forces at cell-matrix adhesions, epithelial junctions, and within the extracellular matrix (ECM), respectively. Insertion sites were selected based on sequence alignments with mammalian orthologs and guided by AlphaFold structural predictions^40^ (**Fig. 4a**; **Extended Data Fig. 6a,b**). For each of these three essential genes, whose loss of function causes embryonic or larval lethality, we tested insertion designs sequentially and stopped at the first design that yielded homozygous viable, fertile, and overtly wild-type animals (**Extended Data Fig. 6c–e**). This required only one design for one gene and two designs for each of the other two (**Extended Data Fig. 6c**), indicating that functional WHaloForce insertions can be obtained at endogenous loci with structure-guided site selection. These lines constitute a *C. elegans* WHaloForce sensor series: CeVin-WF, CeEcad-WF, and CeLam-WF. To generate matched cpHalo controls, we reverted the engineered tryptophan residues in WHalo-Force back to the cpHalo sequence using CRISPR. We also generated C-terminal WHaloForce knock-in controls, CeVin-C-WF, CeEcad-C-WF, and CeLam-C-WF, to place the sensor outside the predicted primary force-bearing geometry of each protein (**Fig. 4a**).

**Figure 4.**
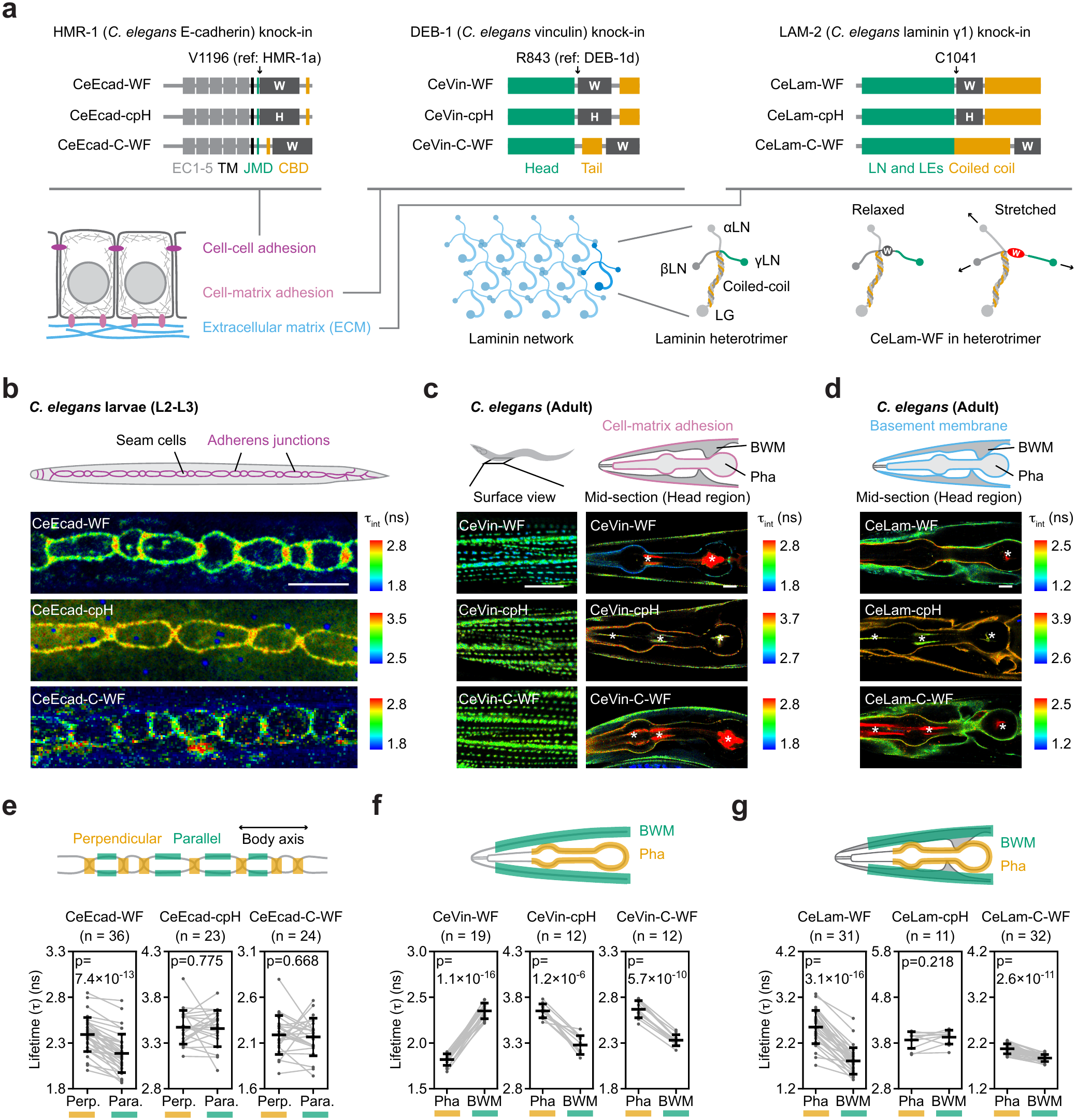
Endogenous WHaloForce knock-ins reveal tissue- and orientation-dependent tension patterns in *C. elegans*. **(a)** Schematics of endogenous WHaloForce, cpHalo, and C-terminal WHaloForce knock-in designs for *C. elegans* E-cadherin/HMR-1, vinculin/DEB-1, and laminin γ1/LAM-2. Schematics of the laminin network and laminin heterotrimer in relaxed and stretched states are also shown. **(b–d)** Schematics of imaged regions and representative FLIM images of endogenous WHaloForce sensors and matched controls in *C. elegans*. **(b)** E-cadherin/HMR-1 sensors and controls at larval epithelial seam-cell junctions (L2-L3). **(c)** Vinculin/DEB-1 sensors and controls in the adult head region. **(d)** Laminin γ1/LAM-2 sensors and controls in the adult head region. Color represents intensity-weighted mean fluorescence lifetime (τ_int_). Color scales were adjusted independently for each image to avoid saturation. Asterisks indicate unbound dye retained in the pharyngeal lumen. **(e–g)** Schematics of regions of interest (ROIs) used for quantification (top) and plots of in-ROI intensity-weighted mean fluorescence lifetime per worm (bottom). **(e)** Orientation-dependent HMR-1/E-cadherin lifetime at seam-cell junctions. Perp. and Para. Indicate perpendicular and parallel directions to the worm body axis, respectively. **(f,g)** Tissue-dependent DEB-1/vinculin **(f)** and LAM-2/laminin γ1 **(g)** lifetimes in the pharynx and body-wall muscle regions. Each point represents one worm. Bars indicate mean ± SD. Statistics: two-sided paired t-tests. All animals were labeled with JFX_554_-HTL. Scale bars, 10 μm.

All nine knock-in lines were homozygous viable, fertile, and overtly wild type, and their protein products localized to the expected adhesion structures or basement membrane ECM (**Fig. 4b–d**; **Extended Data Fig. 6d,e**), indicating that the knock-in fusions retained essential protein function in vivo. As observed with mammalian constructs, C-terminal WHaloForce controls exhibited elevated basal lifetimes relative to internal WHaloForce insertions (**Fig. 4e–g**), indicating that fusion geometry influences intrinsic quenching. We therefore interpreted endogenous WHaloForce measurements using within-construct lifetime contrasts and matched cpHalo and off-axis WHaloForce controls, rather than absolute lifetime values across sensor designs.

Using this framework, FLIM imaging revealed protein-, tissue-, and orientation-dependent lifetime patterns in living worms (**Fig. 4b–g**). We first examined epithelial junctional tension in the larval epidermis, where two rows of seam cells undergo stereotyped divisions while maintaining contacts with neighboring cells. CeEcad-WF showed higher fluorescence lifetime at junctions oriented perpendicular to the worm body axis than at junctions oriented parallel to it, a pattern not observed in the matched controls (**Fig. 4b,e**). This anisotropy indicates that HMR-1/E-cadherin experiences orientation-dependent tension during seam-cell division, consistent with mitotic rounding preferentially loading junctions perpendicular to the division axis.

Next, we examined vinculin and laminin tension in the head region of adult worms. In body-wall muscle, CeVin-WF and controls localized to dense bodies, integrin-based sarcomere attachment structures that connect the contractile apparatus to the muscle membrane and underlying basement membrane^41^ (**Fig. 4c**). CeVin-WF showed lower fluorescence lifetime in the pharynx than in body-wall muscle (Δτ = 0.5 ns; **Fig. 4c,f**), indicating higher tension across DEB-1/vinculin in body-wall muscle. The CeVin-cpH and off-axis CeVin-C-WF controls showed contrasts in the opposite direction (Δτ = -0.4 ns for CeVin-cpH; Δτ = -0.3 ns for CeVin-C-WF; **Fig. 4c,f**), indicating that tissue-dependent effects on cpHalo fluorescence cannot account for the CeVin-WF pattern.

Strikingly, CeLam-WF showed the opposite tissue pattern to CeVin-WF: higher fluorescence lifetime around the phar-ynx than around body-wall muscle (Δτ = 0.7 ns; **Fig. 4d,g**), suggesting increased tensile loading within the laminin network of the pharyngeal basement membrane. Although CeLam-C-WF showed some tissue-dependent lifetime variation, this contrast was much smaller in magnitude than that observed with CeLam-WF (Δτ = 0.2 ns vs. 0.7 ns; **Fig. 4d,g**), indicating that tissue-dependent baseline effects cannot account for the majority of the CeLam-WF pharynx/body-wall difference.

Taken together, these results demonstrate that endogenous WHaloForce knock-ins report molecular tension across multiple force-transducing proteins in living animals. Across cell-matrix adhesions, epithelial junctions, and the basement membrane, WHaloForce revealed protein-, tissue-, and orientation-dependent lifetime patterns consistent with differential molecular tension in vivo. The behavior of the matched cpHalo and C-terminal WHaloForce controls further underscores the importance of interpreting endogenous measurements through within-construct lifetime comparisons and appropriate controls. Beyond enabling in vivo tension measurements, these knock-ins provided experimentally validated insertion geometries that may inform WHaloForce deployment in other animal models.

### A *C. elegans*-guided Vcl-WF knock-in extends WHalo-Force tension imaging to mouse cells and tissues

The *C. elegans* knock-in experiments indicated that endogenous insertion tolerance depends strongly on sensor position. Although the E883 insertion site supported Vin-WF function in cultured mammalian cells and was previously used for a FRET-based vinculin tension sensor^6^, the corresponding insertion site in *C. elegans* DEB-1 (E891) did not yield homozygous viable animals. In contrast, the alternative DEB-1 insertion used for CeVin-WF (R843) produced viable, fertile, and overtly wild-type animals while reporting tissue-dependent vinculin tension in vivo. We therefore generated a mouse vinculin knock-in, *Vcl-WF*, by inserting WHaloForce into the endogenous *Vcl* locus at the site structurally corresponding to the tolerated CeVin-WF insertion (D856), rather than at E883 (**Fig. 5a**; **Extended Data Fig. 6a,b**).

**Figure 5.**
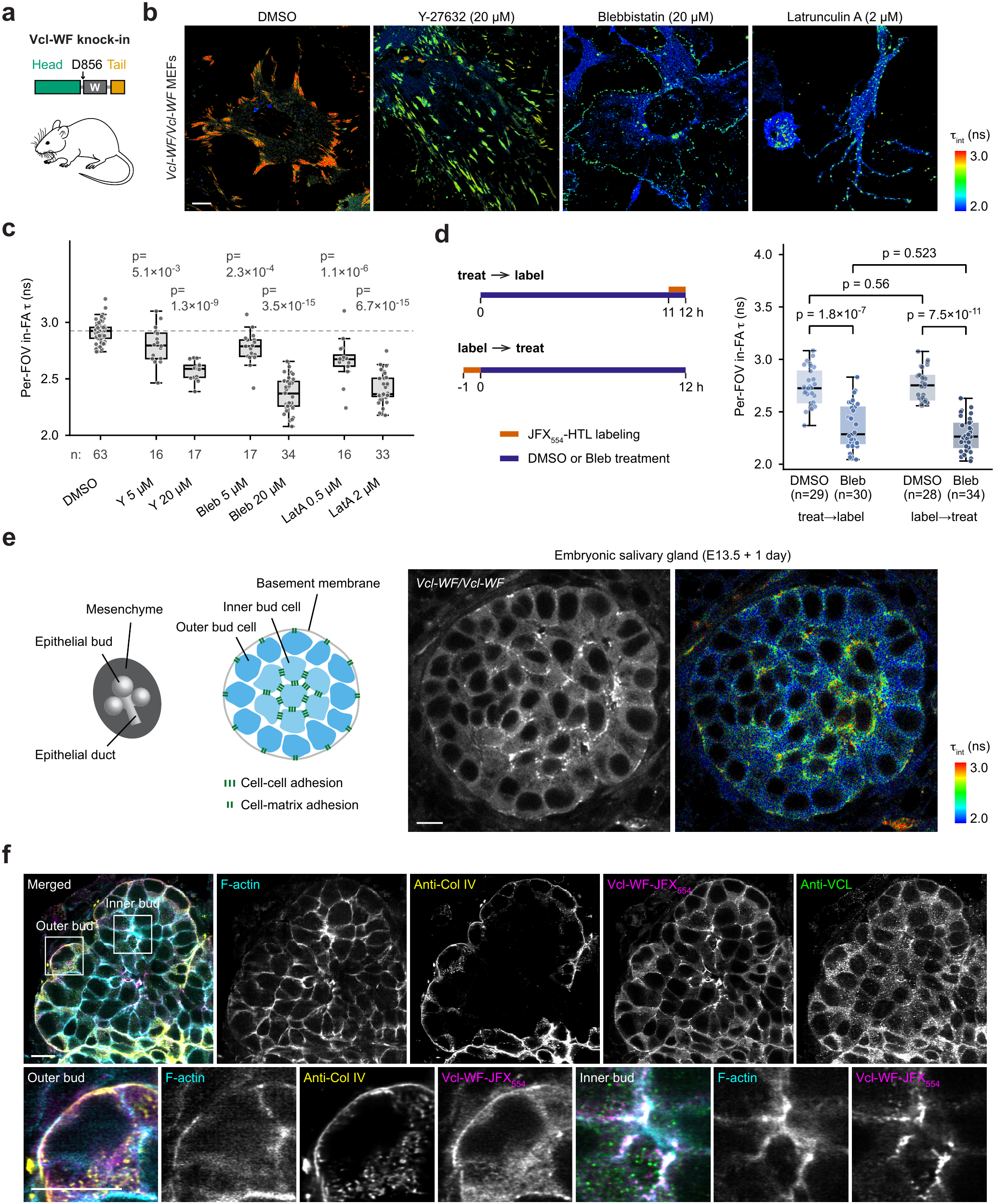
Vinculin tension imaging in cells and tissues from a knock-in mouse model. **(a)** Schematic of the WHaloForce insertion in mouse vinculin, generating Vcl-WF. **(b)** FLIM images of endogenous Vcl-WF at focal adhesions (FAs) in mouse embryonic fibroblasts (MEFs) isolated from homozygous Vcl-WF embryos under the indicated treatment conditions. Cells in **(b,c)** were treated with the indicated drugs and concentrations, or with an equal volume of DMSO as a vehicle control, for 12 h. Cells were labeled with JFX_554_-HTL. Color represents intensity-weighted mean fluorescence lifetime (τ_int_). **(c)** Box-and-scatter plot of per-FOV mean in-FA lifetime under the indicated treatment conditions. Each point represents one FOV; different FOVs contain different cells. Statistics: Mann-Whitney U tests with Holm-Bonferroni correction; reported p values are Holm-adjusted. **(d)** Staged-labeling timeline (left) and box-and-scatter plot of per-FOV mean in-FA lifetime (right). Cells were treated with 20 μM blebbistatin (Bleb) or an equal volume of DMSO for 12 h and labeled with JFX_554_-HTL for 1 h at the end of (treat→label) or before (label→treat) treatment. Each point represents one FOV; different FOVs contain different cells. Data were pooled from two independent experiments. Statistics: Mann-Whitney U tests with Holm-Bonferroni correction; reported p values are Holm-adjusted. **(e)** Schematic of a mouse embryonic salivary gland (left), model of a bud cross section showing inferred cell-cell and cell-matrix adhesion strength (middle), and intensity and FLIM images of a bud cross section from an E13.5 salivary gland cultured for 1 day (right). Salivary glands were labeled with JFX_554_-HTL. Color represents intensity-weighted mean fluorescence lifetime (τ_int_). **(f)** Single-z-section super-resolution images of an E13.5 homozygous Vcl-WF salivary gland cultured for 3 days, labeled with JFX_554_-HTL, and stained with phalloidin (F-actin), anti-collagen IV (Col IV, basement membrane marker), and anti-vinculin (VCL). Scale bars, 10 μm.

Homozygous *Vcl-WF* embryos were recovered from heterozygous intercrosses and appeared morphologically normal at the stage examined, suggesting that this insertion is compatible with essential vinculin function during embryonic development. We also obtained homozygous *Vcl-WF* mice that survived to adulthood and appeared overtly normal. Mouse embryonic fibroblasts (MEFs) derived from *Vcl-WF* embryos showed focal-adhesion-localized fluorescence lifetime patterns (**Fig. 5b**). Inhibition of myosin activity, ROCK signaling, or F-actin polymerization reduced Vcl-WF fluorescence lifetimes at focal adhesions (**Fig. 5c**), indicating that endogenous Vcl-WF responds to actomyosindependent tension in primary mammalian cells.

We next asked whether labeling before or after a sustained reduction in actomyosin activity altered the endogenous reporter response. Blebbistatin produced similar Vcl-WF lifetime decreases under the two labeling schedules for JFX_554_, JF_549_, and JF_571_ (**Fig. 5d; Extended Data Fig. 7**). By comparison, ectopically expressed Vin-WF in NIH-3T3 cells showed labeling-schedule-dependent baseline life-times and variable attenuation of the blebbistatin response across experiments (**Extended Data Fig. 7**). Thus, labeling-history effects varied across the tested vinculin reporter contexts.

We next tested whether endogenous Vcl-WF could be imaged in intact embryonic tissue. In live salivary glands from *Vcl-WF* embryos, Vcl-WF revealed elevated fluorescence lifetimes at adhesions between inner bud epithelial cells and at adhesions between outer bud cells and the basement membrane (**Fig. 5e**). At higher resolution in fixed samples, Vcl-WF partially colocalized with VCL antibody staining and positively correlated with F-actin enrichment (**Fig. 5f**). These patterns support the mechanical heterogeneity we previously proposed in the developing salivary gland, in which combined strong cell-matrix adhesion and weak cell-cell adhesion among outer bud cells drive budding morphogenesis^42^. These results show that dye-labeled endogenous Vcl-WF can be imaged at native expression levels in intact embryonic tissue.

Because matched mouse cpHalo and off-axis WHaloForce knock-in controls have not yet been generated, these tissue data demonstrate endogenous WHaloForce imaging in mammalian organs but do not yet provide a fully controlled, quantitative tension map. Together, these results show that insertion sites identified through endogenous *C. elegans* knock-in screening can guide WHaloForce deployment at mammalian loci, extending FLIM-based molecular tension imaging to primary mouse cells and embryonic tissues.

### WHaloForce identifies ovulation-dependent laminin loading in the *C. elegans* spermathecal basement membrane

We next returned to *C. elegans*, where a single tissue can be followed through repeated cycles of physiological loading across the animal’s lifetime and perturbed genetically. *C. elegans* primarily reproduces as self-fertilizing hermaphrodites. During each ovulation cycle, a mature oocyte is pushed through the spermatheca, expanding its basement membrane by ∼70%^43^. At peak ovulation in day-2 adults, this process repeats more than 100 times per animal within a single day, imposing substantial cyclic mechanical loading on the spermathecal basement membrane. At this stage, CeLam-WF, but not the corresponding cpHalo or C-terminal WHaloForce controls, exhibited higher fluorescence lifetimes at the spermatheca than in neighboring tissues, including the proximal germline and uterus (**Fig. 6a,b**). This local lifetime elevation indicates increased tensile loading within the laminin network at the spermathecal basement membrane. The elevation was absent in L4 animals before ovulation began and became much less pronounced in aged adults after ovulation had ceased (**Fig. 6a,b**), suggesting that increased laminin tension at the spermatheca arises from repeated stretching and recoil during ovulation.

**Figure 6.**
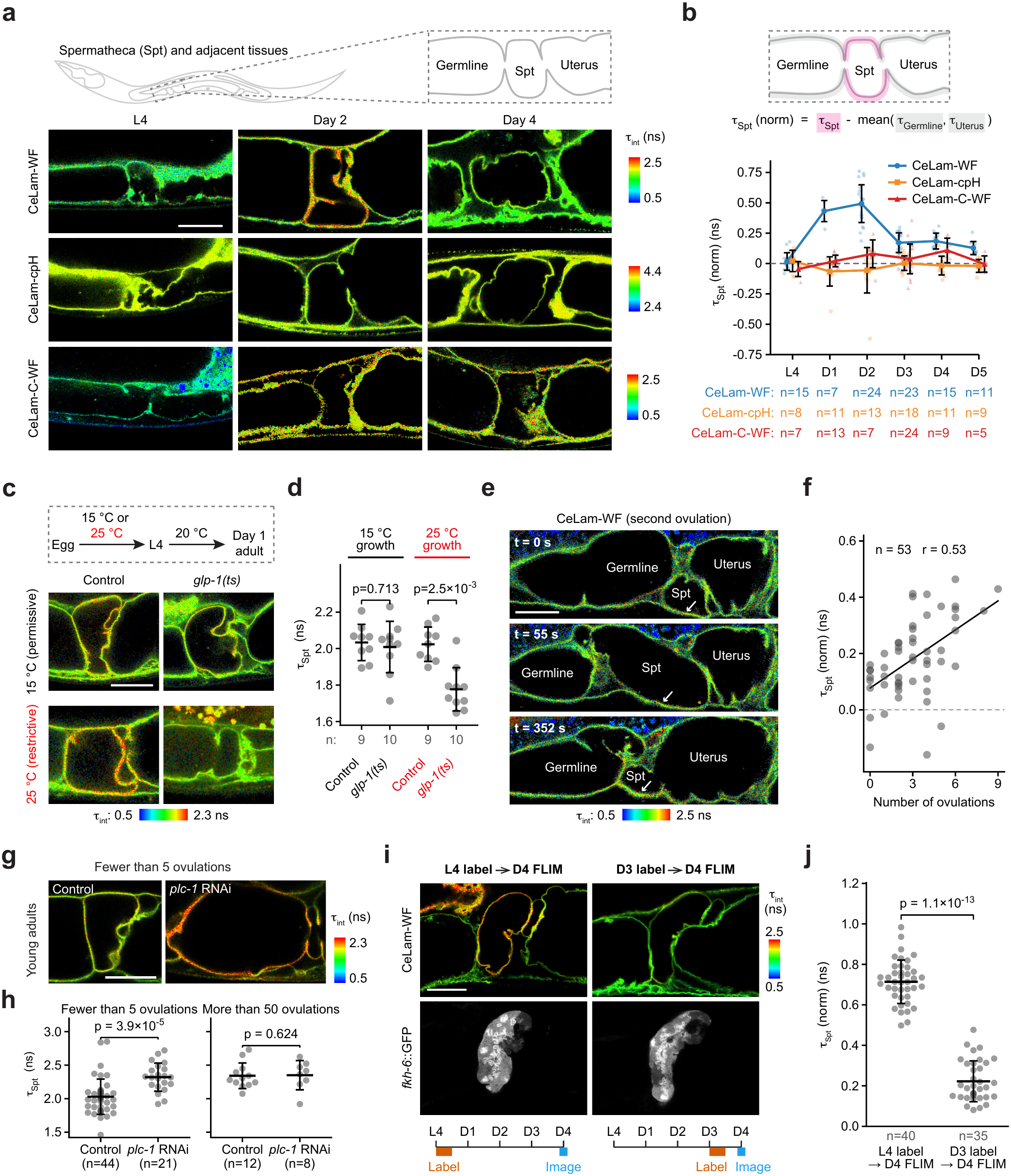
WHaloForce identifies ovulation-dependent laminin loading in the *C. elegans* spermathecal basement membrane. **(a)** Schematics of the imaged region and representative FLIM images of the indicated ages and genotypes. Color represents intensity-weighted mean fluorescence lifetime (τ_int_). Color scales were adjusted independently for each image to avoid saturation. **(b)** Schematics of regions of interest (ROIs) used for quantification (top) and plot of local lifetime contrast at the spermathecal basement membrane over age (bottom). Local lifetime contrast, or τ_Spt_ (norm), was defined as the intensity-weighted mean fluorescence lifetime in the spermathecal ROI minus the average lifetime in the two neighboring tissue ROIs, the proximal germline and uterus. Each point represents one worm; error bars indicate SD; n, worms per condition. Two-sided Mann-Whitney U tests with Holm-Bonferroni correction. At L4, CeLam-WF did not differ from either control (adjusted p = 0.51 versus CeLam-cpH; p = 0.19 versus CeLam-C-WF). From day 1 to day 5, CeLam-WF differed from both controls (adjusted p ≤ 0.035), except from CeLam-C-WF at day 4 (p = 0.19). CeLam-WF τ_Spt_ (norm) was higher at day 2 than at L4 and at days 3 to 5 (adjusted p ≤ 8.4 × 10^-6^), but did not differ from day 1 (p = 0.32). Full statistics are provided in Source Data. **(c)** Schematic of temperature shifting (top) and FLIM images (bottom) of control and *glp-1(ts)* mutant worms expressing CeLam-WF, raised at the permissive temperature (15 °C) or restrictive temperature (25 °C) until the L4 stage and imaged as day-1 adults. Color represents intensity-weighted mean fluorescence lifetime (τ_int_). **(d)** Plot of intensity-weighted mean fluorescence lifetime of CeLam-WF at the spermathecal basement membrane under the indicated conditions. Statistics: Mann-Whitney U tests with Holm-Bonferroni correction; reported p values are Holm-adjusted. **(e)** FLIM images of CeLam-WF before oocyte entry, during oocyte passage, and after oocyte exit during an early ovulation event (the second ovulation). Arrows point to the spermathecal basement membrane. Color represents intensity-weighted mean fluorescence lifetime (τ_int_). **(f)** Scatter plot of local lifetime contrast at the spermathecal basement membrane, calculated as defined in (b), versus the cumulative number of ovulations experienced by the corresponding spermatheca in young adults. Each dot represents one spermatheca; n = 53 spermathecae from 42 worms. Line, linear fit; Pearson r = 0.53 (95% confidence interval, 0.28–0.71). The confidence interval was calculated using 50,000 whole-worm bootstrap resamples, retaining all spermathecal measurements from each sampled worm together. **(g)** FLIM images of CeLam-WF under control or *plc-1* RNAi conditions in young adults with fewer than 5 ovulations per spermatheca. Color represents intensity-weighted mean fluorescence lifetime (τ_int_). **(h)** Plot of intensity-weighted mean fluorescence lifetime of CeLam-WF at the spermathecal basement membrane under the indicated conditions. n, worms per condition. Ovulation counts are per spermatheca. Statistics: Mann-Whitney U tests with Holm-Bonferroni correction; reported p values are Holm-adjusted. **(i)** FLIM images of CeLam-WF (top), fluorescence images of the spermathecal marker *fkh-6::GFP* (middle), and labeling timelines (bottom) for worms labeled with JFX_554_-HTL at L4 or on adult day 3 and imaged on adult day 4. Labeling and imaging times are marked on the timelines. Color represents intensity-weighted mean fluorescence lifetime (τ_int_). **(j)** Plot of local lifetime contrast at the spermathecal basement membrane, calculated as defined in (b), under the indicated labeling schedules. Each point represents one worm; bars indicate mean ± SD; n, worms pooled from two independent experiments. Statistics: two-sided Mann-Whitney U test. All animals were labeled with JFX_554_-HTL. Scale bars, 20 μm.

To test whether ovulation is required for this increase in laminin tension, we crossed CeLam-WF into the temperature-sensitive *glp-1(e2144)* mutant background^44^, in which germline function is disrupted at the restrictive temperature (25 °C). When grown at 25 °C until the L4 stage, *glp-1(ts)* adults failed to ovulate. Under these conditions, CeLam-WF fluorescence lifetime at the spermathecal basement membrane remained low in day-1 adults, an age at which control animals already showed elevated laminin tension (**Fig. 6c,d**). Thus, elevated laminin tension at the spermatheca depends on ovulation.

Surprisingly, at the peak ovulation stage, CeLam-WF fluorescence lifetime remained elevated at the spermatheca even after the fertilized oocyte exited and the basement membrane returned toward its relaxed geometry (**Fig. 6a**). To examine the initial response, we performed live FLIM imaging during early ovulation events in young adults. Over a single ovulation event, CeLam-WF fluorescence lifetime showed only a modest increase (**Fig. 6e**; **Video S4**). Furthermore, CeLam-WF lifetime at the spermathecal basement membrane positively correlated with the number of ovulations per spermatheca in young adults (**Fig. 6f**; see Methods).

To test how increasing the extent and duration of basement membrane stretching affects the laminin response, we performed *plc-1* RNAi, which impairs spermathecal contraction and oocyte exit, causing prolonged and enhanced spermathecal distension as multiple oocytes accumulate within the spermatheca^45,46^. Strikingly, in young adults subjected to *plc-1* RNAi, CeLam-WF fluorescence lifetime at the spermathecal basement membrane reached levels comparable to controls that had undergone more than 50 ovulations after only a few ovulation events (**Fig. 6g,h**). Together with the matched sensor controls and the loss of the local life-time elevation when ovulation was prevented, these results support a tension-dependent origin of the spermathecal

#### CeLam-WF contrast

To examine whether labeling history shapes this tension-dependent response, we compared labeling schedules at a common adult-day-4 endpoint. Worms labeled overnight starting at L4, before the onset of ovulation, and subsequently maintained on dye-free plates showed higher local CeLam-WF lifetime contrast at the spermatheca than agematched worms labeled overnight on adult day 3 (**Fig. 6i,j**). Thus, laminin remaining from the early-labeled population shows a stronger CeLam-WF response after several days of ovulatory activity than the population labeled later in adulthood.

Together, these experiments show that the laminin network bears load during physiological ovulation, linking deformation of the spermatheca to molecular force transmission through its basement membrane.

## Discussion

In this study, we introduce WHaloForce, a compact chemigenetic molecular tension sensor that converts mechanical force into a change in dye fluorescence lifetime. Building on PET-based HaloTag engineering developed for fluorescence lifetime multiplexing^47^ and calcium sensing^16^, WHaloForce extends tryptophan-rhodamine PET quenching from chemical to mechanical sensing, establishing, to our knowledge, the first chemigenetic PET-based tension sensor. Single-molecule measurements showed a switch-like unquenching transition near 5 pN, and WHalo-Force imaging resolved beat-coupled lifetime oscillations in cardiomyocytes at ∼1.2 Hz. When inserted into force-bearing proteins, WHaloForce reported dynamic, myosin-dependent vinculin tension at focal adhesions, anisotropic tension at epithelial junctions, opposite vinculin and laminin tension patterns between tissues in living animals, and ovulation-dependent laminin loading in the spermathecal basement membrane.

WHaloForce complements rather than replaces existing molecular tension sensors. FRET-based sensors remain the most established genetically encoded approach, reporting force through energy transfer across an extensible linker between two fluorescent proteins, read out ratiometrically or by donor fluorescence lifetime^6,8–11,13^. Recruitment-based sensors convert force-dependent exposure of a cryptic motif into the intensity of a bound partner^15^, and cpEGFP-based indicators report deformation through intensity normalized to a reference fluorophore^48^. FRET and cpEGFP readouts are analog, varying continuously with the load each molecule carries; recruitment-based readouts are threshold-like, but report the binding of a separate partner rather than the state of the sensor itself. WHaloForce differs from these in two respects. First, the sensing element is a single HaloTag domain with one bound dye, requiring no donor-acceptor pair, no recruited partner, and no reference channel. It is correspondingly compact and occupies a single spectral channel. Most force measurements here used JFX_554_, whose brightness enables use of the sensor at endogenous expression levels, essential for the knock-in experiments; the staged-labeling experiments additionally used JF_549_ and JF_571_. Because the readout is a fluorescence lifetime rather than an intensity, it also remains quantitative where labeling density, expression level, and imaging depth vary, a property shared with lifetime-based FRET but not with intensity-based designs.

Second, WHaloForce is digital at the level of the individual molecule. cpWHalo occupies one of two discrete quenching states, so each sensor is either quenched or unquenched, and switching rates depend steeply on load: unquenching was 12-fold faster at 6 pN and 28-fold faster at 8 pN than at 5 pN, whereas re-quenching kinetics remain incompletely defined. Analog and digital sensors are both read out as an average over many molecules per pixel, but they average different quantities. An analog sensor averages a continuously varying force-dependent signal, so a narrow and a broad force distribution can give the same signal. A digital sensor averages state occupancy, and so responds to the fraction of molecules that have crossed the transition^15,49^. WHaloForce therefore reports where tension drives switch-like unquenching rather than absolute force magnitude.

Because occupancy is kinetically set, WHaloForce is a kinetic tension sensor rather than an equilibrium force gauge. When unquenching and re-quenching are both fast relative to the process under study, occupancy tracks changing load; slower re-quenching can prolong the response after unloading. The temporal response therefore depends on the reporter context and need not be instantaneous.

The staged-labeling experiments illustrate this context dependence. Endogenous Vcl-WF in MEFs showed substantial blebbistatin responses under both schedules, with no consistent attenuation by prelabeling across three dyes and two experiments. Ectopically expressed Vin-WF in NIH-3T3 cells showed labeling-schedule-dependent baselines and a variable drug-by-schedule interaction. Expression regime, cell background, and insertion geometry may contribute to this difference, but their effects are confounded. For comparisons of sustained conditions, treatment, labeling, washout, and imaging intervals should therefore be standardized.

WHaloForce identified ovulation-dependent loading of the spermathecal laminin network. The persistent response in laminin remaining from the early-labeled population could reflect retained or recurrent loading, slow reporter recovery, selective retention of loaded laminin, or labeling and accessibility effects. If residual laminin tension contributes, fibulin-collagen remodeling^43^ and cell-matrix attachment could sustain or redistribute the load.

WHaloForce also revealed tissue-specific load routing. In the head, CeVin-WF lifetime was lower in the pharynx than in body-wall muscle, whereas CeLam-WF showed the opposite pattern: vinculin tension was higher in body-wall muscle, and laminin tension higher in the pharynx (**Fig. 4c,d**). Adhesion and matrix tension therefore run in opposite directions between these two adjacent tissues, rather than rising and falling together as they would if one tissue were simply under greater load.

Body-wall muscle transmits contractile force through integrin-containing dense bodies to the basement membrane, hypodermis, and cuticle^41^, consistent with higher DEB-1/vinculin tension there. The pharynx, in contrast, is an enclosed muscular pump isolated by its own basement membrane^50^, which is roughly twice as thick as those around most other *C. elegans* tissues (∼45 nm compared with ∼20 nm)^51,52^. Higher CeLam-WF lifetime there may reflect loading of that reinforced shell as it resists organ deformation and the repeated forces of feeding. The same tissue-scale deformation may therefore load different molecular elements to different extents, depending on boundary conditions, curvature, and matrix architecture.

Interpreting WHaloForce measurements requires care on two related points. First, lifetime depends on sensor geometry: the different basal lifetimes of C-terminal fusions in cultured cells and *C. elegans* (**Figs. 3c,f and 4e–g**) indicate that fusion orientation and local protein context alter quenching efficiency, as expected for a mechanism that depends on the geometry between the engineered tryptophan and the bound dye. Geometry also sets how load reaches the sensor: in the optical trap, force was applied between the termini through DNA handles, which need not match the load path at a given insertion site in cells. Second, CeVin-cpH showed a tissue-dependent lifetime difference opposite in sign to CeVin-WF (**Fig. 4c,f**), which cannot reflect the intended mechanism because cpHalo lacks the quenching tryptophan; it more likely reflects effects of the local environment on the dye, such as pH, polarity, viscosity, or oxidative state. Absolute lifetimes should therefore not be compared across host proteins, insertion sites, or fusion geometries; lifetime differences should be assigned to tension only when supported by matched cpHalo controls, off-axis controls where feasible, and force perturbations. WHaloForce is best used for relative measurements across space and time.

Two practical constraints apply. Because the core of the sensor is HaloTag, WHaloForce depends on efficient dye labeling. Dye delivery, tissue penetration, ligand washout, and background fluorescence vary across organisms and tissues, which will require optimization. The sensor also requires enough photons for robust lifetime estimation, which matters most for endogenous knock-ins where expression is fixed and often low, and lifetime fitting remains sensitive to background and model choice. Systematic screening of circular permutation sites, quenching residues, dye chemistries, and linker architectures should yield variants with distinct thresholds, expanded dynamic range, and improved environmental robustness. Even in its present form, a single sensor design reported tension on four forcebearing proteins in cultured cells, mouse tissue, and living *C. elegans*, spanning single molecules to intact tissue, and we expect WHaloForce to be broadly useful for FLIM-based molecular tension imaging across diverse proteins, tissues, and organisms.

## Acknowledgements

We thank the Molecular Genomics, Viral Tools, Flow Cytometry, Janelia Experimental Technology (jET), Gene Targeting & Transgenics, Vivarium, Project Technical Resources (PTR), Anatomy and Histology, Invertebrate Shared Resource, and Open Chemistry at Janelia Research Campus for support. We especially thank C. Huang, A. Ludlow, H. A. Yi, C. Li, and J. Arnold. We thank M. Park and M. Wang for help with *C. elegans* RNAi experiments; D.K. Cheerambathur (U. Edinburgh) for sharing *C. elegans* CRISPR knock-in protocols; C. de Caestecker, A. Desai, Z. J. Gartner, R. Vale, M. Wang, and K. M. Yamada for critical comments on the manuscript. Some *C. elegans* strains were provided by the CGC, which is funded by NIH Office of Research Infrastructure Programs (P40 OD010440). This work was supported by the Howard Hughes Medical Institute. The contributions of J.L., S.P., and T.H. are additionally supported by the US National Institutes of Health (R35 GM122569). The contributions of R.J.B., Q.Y., and H.W. are supported by the US National Institutes of Health (R01 HL179359).

## Author Contributions

Conceptualization: D.W., E.R.S., S.W.

Methodology: D.W., E.A.M., J.L., S.P., H.F., X.L., H.S., H.W., A.G.T., L.D.L., E.R.S., T.H., S.W.

Software: D.W., J.L., S.P., S.W.

Validation: D.W., E.A.M.

Formal analysis: D.W., J.L., S.P., S.W.

Investigation: D.W., E.A.M., J.L., S.P., H.F., K.A.H., R.J.B., Q.Y., X.L., S.W.

Resources: H.F., E.R.S., A.G.T., L.D.L., R.J.B., Q.Y., H.W.

Data curation: D.W., J.L., S.P., S.W.

Visualization: D.W., S.W.

Supervision: H.S., H.W., A.G.T., L.D.L., E.R.S., T.H., S.W.

Funding acquisition: H.S., H.W., A.G.T., L.D.L., E.R.S., T.H., S.W.

Writing - original draft: D.W., S.W. Writing - review & editing: All authors.

## Methods

### Use of generative AI tools

During manuscript preparation, OpenAI ChatGPT (5.5, 5.6-sol) and Anthropic Claude (Opus 4.8, Opus 5) were used for language editing. OpenAI Codex (v0.142.5; model: gpt-5.5, 5.6-sol) and Anthropic Claude Code (v2.1.220; model: Opus 4.8) were used to assist with code drafting, debugging and documentation. The authors reviewed and validated all text and code outputs, verified the analyses and take responsibility for the final content.

### Mouse work

All mouse experiments were approved by the Janelia Research Campus Institutional Animal Care and Use Committee (IACUC) under protocol 25-0279. Mice were maintained on an FVB/N genetic background. Embryonic day 0.5 (E0.5) was defined as noon on the day a vaginal plug was detected.

### *C. elegans* maintenance

The *C. elegans* strains used in this study are listed in **Table S1**. Unless otherwise indicated, worms were maintained at 20 °C on standard nematode growth medium (NGM) plates seeded with OP50 bacteria. For feeding RNAi experiments, worms were maintained on NGM plates seeded with HT115 bacteria carrying either the empty L4440 vector or a *plc-1* RNAi construct. Worms carrying the temperature-sensitive *glp-1(e2144)* allele were maintained at the permissive temperature of 15 °C. To disrupt *glp-1* function during germline development, worms were shifted from 15 °C to the restrictive temperature of 25 °C during early embryogenesis, maintained at 25 °C until the L4 stage, and returned to 20 °C thereafter.

### Culture of immortalized cell lines

HEK293T cells were obtained from Takara Bio (632180), and NIH-3T3 and MD-CKII cells were obtained from the laboratory of Kenneth Yamada at the National Institutes of Health. HEK293T and MDCKII cells were cultured in DMEM (Thermo Fisher, 10566024) supplemented with 10% fetal bovine serum (FBS; Cytiva, SH30071.03 or Thermo Fisher, 26140079) and penicillin-streptomycin (Thermo Fisher, 15140122). NIH-3T3 cells were cultured in DMEM supplemented with 10% bovine calf serum (BCS; ATCC, 30-2030) and penicillin-streptomycin. Cells were maintained at 37 °C with 5% CO_2_, passaged using trypsin-EDTA (Thermo Fisher, 25300120 or 25200114), and used within 20 passages.

### Isolation and culture of mouse embryonic fibroblasts

Mouse embryonic fibroblasts (MEFs) were isolated from E13.5 embryos. For *Vcl-WF* embryos, a tail tissue sample from each embryo was used for genotyping, and only homozygous embryos were used for salivary gland culture and MEF isolation. After removal of the head and internal organs, bodies from up to three embryos were pooled in a 10-cm dish containing approximately 8 mL of prewarmed 0.25% trypsin-EDTA (Thermo Fisher Scientific, 25200114) and minced thoroughly using microsurgical scissors, forceps, and a scalpel. The tissue was triturated several times with a 10-mL serological pipette and incubated at 37 °C for 10 min. The suspension was then triturated with a 5-mL pipette, incubated at 37 °C for an additional 10 min, and triturated again with a 10-mL pipette. The digest was transferred to a 50-mL tube, mixed with 20 mL of prewarmed complete MEF medium, consisting of DMEM supplemented with 10% FBS and penicillin-streptomycin, and allowed to settle for approximately 5 min. The cell suspension was transferred to a T75 flask without disturbing the sedimented tissue fragments, supplemented with an additional 10 mL of complete MEF medium, and maintained at 37 °C with 5% CO_2_. MEFs were used within 10 passages.

### hiPSC origin and maintenance

Human induced pluripotent stem cells (hiPSCs) were originally derived from a healthy male donor of Asian ancestry with informed consent under a University of Pittsburgh IRB-approved protocol (CR21100125-004). hiPSCs were maintained on Matrigelcoated tissue-culture plates. Plates were coated with Matrigel (Corning, 354277) diluted 1:250 in DMEM/F12 1:1 (Cytiva, SH30023.01) corresponding to ∼40 μg/mL. Cells were cultured in Essential 8 medium (E8; Thermo Fisher, A1517001), which was replaced daily, and maintained in a humidified incubator at 37 °C with 5% CO_2_ and 5% O_2_. Cultures were passaged every 4–5 days at ∼70–80% confluence using either 0.5 mM EDTA in PBS or Accutase solution (Sigma A6964). Y-27632 (Selleckchem, S1049) was added at 10 μM during passaging to enhance cell survival and attachment and was removed the following day by replacing the medium with fresh E8. Colony morphology was monitored routinely, and overgrowth was avoided to maintain healthy, undifferentiated cultures.

### iPSC-derived cardiomyocyte differentiation

Human iP-SCs were differentiated into cardiomyocytes using small-molecule Wnt modulation. Differentiation was initiated when hiPSC cultures reached ∼85% confluence. On day 0, cultures were washed with RPMI 1640 (Gibco, 11875093) and treated with RPMI 1640 supplemented with B27 minus insulin (1:50 supplement to medium) (Thermo Fisher; A1895601) and 6 μM CHIR99021 (Selleckchem, S2924) for 2 days. On day 2, the medium was replaced with RPMI 1640 supplemented with B27 minus insulin without CHIR99021. On day 3, cells were treated with RPMI 1640 supplemented with B27 minus insulin and 5 μM IWR-1-endo (Selleckchem, S7086) for 2 days. On day 5, the medium was replaced with RPMI 1640 supplemented with B27 minus insulin. On day 7, cells were switched to RPMI 1640 supplemented with complete B27 (Thermo Fisher; 17504044). After spontaneous contractions were observed, typically by days 9–10, cells were treated with glucose-free RPMI 1640 (Gibco, 11879020) supplemented with B27 for 2 days, then maintained with RPMI 1640 supplemented with B27. No antibiotics were used during any of the maintenance and differentiation steps.

### iPSC-derived cardiomyocyte maintenance and lentiviral transduction

Differentiated iPSC-derived cardiomy-ocytes (iCMs) were maintained on Matrigel-coated surfaces. Surfaces were coated with Matrigel diluted 1:250 from a ∼10 mg/mL stock in DMEM/F-12 and incubated for at least 1 h at 37 °C before use. iCMs were maintained in RPMI 1640 supplemented with B27 and passaged using TrypLE 10X (dilute to 1X with HBSS supplemented with 1 mM EDTA for usage; Gibco, A1217701) or TrypLE Express Enzyme (1X) (Thermo Fisher, 12604013). For the first 1–2 days after thawing frozen iCMs, cells were maintained in the recovery medium consisting of RPMI 1640 supplemented with B27, 10% heat-inactivated FBS (Thermo Fisher, 26140079), 2 μM CHIR99021, and 10 μM Y-27632. For lentiviral transduction, iCMs were incubated with virus in the presence of 8 μg/mL polybrene (Millipore-Sigma, H9268) at an MOI of 80 in a 24-well plate for 8–16 h and then replated into 8-well coverslip-bottom imaging chamber slides (ibidi, 80807).

### Lifetime-beating coupling analysis in cardiomyocytes

Spontaneously beating iCM fields were treated as independent experimental units. The experiment was repeated in two independent transductions with similar results; the dataset shown comprised 27 Vin-WF and 23 Vin-cpH fields, each containing a single cell. Time series of 100–200 frames (128 × 128 pixels; 0.286 μm per pixel) were acquired at a median interval of 0.167 s (∼6 Hz; Nyquist frequency, 3 Hz), with recorded timestamps used for all frequency and lag calculations. Focal adhesions were segmented independently in each frame using a deterministic rolling-ball plus adaptive-threshold pipeline applied identically to every frame (parameters in the analysis code). The union of per-frame masks defined the fixed in-FA region; its brightest 25% of pixels, determined from timeaveraged photon counts, defined the bright-FA region containing mature costameres. For each frame, photon arrivals within each region were pooled and fit by non-negative least squares against the reconvolved instrument response to obtain τ_int_(t). Pooling before fitting avoids the shared photon-count fluctuations that couple per-pixel intensity-weighted lifetimes to motion, which can generate apparent coupling in the force-insensitive control. Beating was represented by the mean absolute frame-to-frame intensity difference within the in-FA region, with f_0_ defined as its dominant spectral peak. Power spectra of lifetime and motion were computed as Lomb-Scargle periodograms (a least-squares generalization of the Fourier periodogram for unevenly sampled data) evaluated at the recorded frame timestamps. Coupling was calculated after zero-phase, second-order Butterworth filtering over f_0_ ± 50%, as the maximum cross-correlation within ±1/2 beat cycle, with sub-frame lag estimated by parabolic interpolation of the cross-correlation peak. Reported values are from the whole-field bright-FA region; statistical analysis is described under Statistics.

### WHaloForce design and molecular constructs

WHalo-Force, cpHalo, and terminal WHaloForce control constructs were generated in the positions and configurations shown in the corresponding figure schematics. DNA constructs were either synthesized by Twist Bioscience or assembled using Gibson Assembly. All plasmids were verified by whole-plasmid sequencing performed by Plasmidsaurus or GENEWIZ. Complete plasmid sequences have been deposited with Addgene (IDs #261833–#261846). Endogenous WHaloForce, cpHalo, and C-terminal WHaloForce knock-ins were generated at the loci shown in the corresponding figure schematics. The *C. elegans* strains used in this study are listed in **Table S1**, genotyping primers for *C. elegans* and mouse knock-ins in **Table S2**, and guide RNA and homology-directed repair (HDR) donor sequences in **Table S3**.

### Generation of knock-in *C. elegans* strains

Endogenous knock-ins were generated using a modified Cas9 ribonucle-oprotein co-CRISPR protocol based on Paix et al. (2015)^53^. Gene-specific and *dpy-10* crRNAs were combined at a 3:1 molar ratio, duplexed with tracrRNA, and complexed with Cas9 protein. Linear double-stranded DNA repair templates were amplified using Q5 DNA polymerase (NEB), purified using a QIAquick PCR Purification Kit (Qiagen), and concentrated by heat block evaporation followed by air dry. A typical injection mixture contained 2.5 μM Cas9, 3 μM total duplexed RNA, 1 μM *dpy-10* repair oligonucleotide, and the concentrated gene-specific repair template (2-4 μM). The mixture was injected into the germline of young adult hermaphrodites. Roller F1 progeny were screened by PCR, and candidate edits were confirmed by Sanger sequencing. The *dpy-10* mutation was subsequently removed by selecting non-roller progeny. The resulting strains and editing reagents are listed in **Tables S1** and **S3**, respectively.

### Generation of knock-in mice

Knock-in mice were generated on an FVB/N genetic background using a modified 2C-HR-CRISPR method^54^. Cas9-mSA mRNA (75 ng/μL), sgRNA (50 ng/μL), and biotinylated PCR donor DNA (20 ng/μL) were microinjected into the cytoplasm of two-cell embryos, which were transferred immediately to pseudopregnant recipients. Cas9-mSA mRNA was synthesized from pCS2+Cas9-mSA (Addgene, 103882), and the sgRNA was synthesized by Integrated DNA Technologies (IDT). Donor DNA was amplified using Q5 High-Fidelity DNA Polymerase from a plasmid synthesized by Twist Bioscience using biotinylated primers (IDT) and purified using a Qiagen PCR purification kit without gel extraction. The *Vcl-WF* donor contained left and right homology arms of 1,117 and 1,244 bp, respectively. Founders were screened by PCR and confirmed by whole-amplicon sequencing using primers outside the homology arms. Genotyping primers and editing reagents are listed in **Tables S2** and **S3**, respectively.

### Salivary gland isolation, culture, mounting, and labeling

Submandibular salivary glands were isolated, cultured, and mounted as described previously^42^. Briefly, gland explants were dissected from E13.5 embryos and cultured at 37 °C with 5% CO_2_ on a polycarbonate filter (Cytiva, 10417001) floating on organ culture medium: DMEM/F-12 supplemented with penicillin-streptomycin (1X), 150 μg/mL vitamin C (MilliporeSigma, A7506), and 50 μg/mL transferrin (MilliporeSigma, T8158). For labeling, glands were cultured overnight in organ culture medium containing 100 nM JFX_554_-HTL, then washed three times in dye-free medium, 1–2 h per wash. For imaging, a 0.12-mm double-adhesive imaging spacer with a 9-mm opening (Grace Bio-Labs, 654002) was attached to the coverslip bottom of a 35-mm imaging dish (ibidi, 81158). The filter carrying the glands was inverted and attached to the spacer, sandwiching the glands between the filter and the coverslip. The spacer set the gap and prevented the glands from being crushed.

### Salivary gland immunostaining and cryo-sectioning

Salivary glands were fixed overnight at 4 °C in 4% PFA (EMS, 15710), permeabilized for 30 min in PBSTx (PBS with 0.2% Triton X-100), blocked for 1 h in 5% donkey serum in PBSTx, and incubated with primary antibodies for 3 days at 4 °C. Glands were washed four times in PB-STx, at least 15 min per wash, incubated with secondary antibodies and Alexa Fluor 405 Plus phalloidin (Thermo Fisher, A30104; 1:400) for 3 days at 4 °C, washed four more times in PBSTx as above, and mounted for imaging. Primary antibodies were anti-vinculin (MilliporeSigma, V9131; 1:250) and anti-collagen IV (MilliporeSigma, AB769; 1:200). Secondary antibodies were Alexa Fluor 488 donkey antigoat (Jackson ImmunoResearch, 705-546-147; 1:200) and Alexa Fluor 647 donkey anti-mouse (Jackson ImmunoResearch, 715-606-151; 1:200). All incubations were at room temperature with gentle rocking unless otherwise noted. For super-resolution microscopy, samples were post-fixed overnight at 4 °C in 4% PFA, rinsed in PBS, embedded in a cryomold filled with OCT medium, and cryo-sectioned; sections were mounted directly onto the center of a No. 1.5 coverslip.

### Lentivirus production and titration

Lentivirus was produced either in-house or by the Janelia Research Campus Viral Tools Team. In both workflows, 293-derived packaging cells were co-transfected with the lentiviral transfer plasmid and the packaging plasmids psPAX2 (Addgene, 12260) and pMD2.G (Addgene, 12259). For in-house production, HEK293T cells were transfected by calcium phosphate precipitation in the presence of 25 μM chloroquine. Viral supernatants were collected on two successive days, pooled, filtered through a 0.45-μm filter, concentrated by overnight PEG (System Biosciences, LV825A-1) precipitation followed by centrifugation at 1,500 × g for 30 min at 4 °C, resuspended in DMEM/F-12 (Thermo Fisher, 11039047), and stored at ™80 °C. Viral titers were estimated using Lenti-X GoStix Plus (Takara Bio, 631281). For production by the Viral Tools Team, HEK293T cells were transfected using PEI MAX (Kyforabio, 24765-1), and viral supernatants were collected at 48 and 72 h, filtered through a 0.22-μm filter, concentrated using 100-kDa molecular-weight-cutoff centrifugal filters (MilliporeSigma, UFC710008), and purified by ultracentrifugation through a 20% sucrose cushion. Viral titers were determined by digital PCR. Concentrated viral stocks were resuspended in DMEM/F-12 and stored at ™80 °C.

### Lentiviral transduction of immortalized cell lines

All experiments using immortalized cell lines were performed with stably transduced cells. Subconfluent HEK293T, NIH-3T3, and MDCKII cultures were transduced at a multiplicity of infection (MOI) of 10–100 in the presence of 8 μg/mL polybrene (MilliporeSigma, H9268) for 8–16 h, after which the virus-containing medium was replaced with fresh culture medium. Transduced cells were expanded and used directly when transduction was sufficiently efficient and uniform. Cultures with low or heterogeneous transduction were enriched by flow cytometric sorting. In selected cases, clonal lines were established by limiting dilution and selected for normal cell morphology and consistent construct expression patterns.

### HaloTag labeling of cultured cells

Unless otherwise indicated, cultured cells were labeled with JFX_554_-HaloTag ligand (JFX_554_-HTL). Cells were incubated with 100 nM dye-HTL in the corresponding culture medium for 30–60 min and then washed three times for 5 min each with prewarmed medium. For the dye-screening experiment in HEK293T cells, the indicated dye-HTLs were used in place of JFX_554_-HTL. For imaging of HEK293T, NIH-3T3, and MEF cells, coverslip-bottom imaging chambers (ibidi) were coated with 10 μg/mL fibronectin (MilliporeSigma, FC010) in PBS for at least 30 min at room temperature or 37 °C to promote cell attachment.

### Drug treatments

NIH-3T3 and MEF cells were treated with blebbistatin, Y-27632, or latrunculin A at the concentrations indicated in the corresponding figures and legends, or with an equal volume of DMSO as a vehicle control, for 12 h before imaging. Except for prelabeling in the label→treat experiments, drugs were present throughout labeling and imaging. Vendors and catalog numbers: blebbistatin (MilliporeSigma, 203391), Y-27632 (MilliporeSigma, Y0503), latrunculin A (MilliporeSigma, 428026).

### Cell spreading assay with ROCK inhibition

NIH-3T3 cells were trypsinized, resuspended in culture medium at 1 × 10^5^ cells/mL, and mixed 1:1 with 200 nM JFX_554_-HTL to give 5 × 10^4^ cells/mL and 100 nM dye-HTL. Y-27632 (5 μM final, 1:2,000 from a 10 mM stock; MilliporeSigma, Y0503) or an equal volume of DMSO was added, and cells were labeled for 30 min in a 37 °C water bath, washed once, and resuspended at 5 × 10^4^ cells/mL in fresh medium containing 5 μM Y-27632 or DMSO. Cells were seeded at 300 μL per well and FLIM acquisition was started immediately after positions were identified. Six to seven positions were imaged every 5 min for 125 min (26 frames) at 512 × 512 pixels and 3X zoom.

### Uniaxial stretching assay

PDMS membranes (Specialty Manufacturing Inc., gloss silicone sheeting, thickness 0.005 in / 127 μm) were cut and mounted in a custom 3D-printed uniaxial stretching chamber (jET, Janelia Research Campus) adapted from a published design^39^ (**Extended Data Fig. 5a**). The device was calibrated by marking two reference dots on a mounted membrane and measuring interdot distance as a function of applied turns of the knob; linear regression gave 1.14 mm per turn, corresponding to ∼5.7% strain per turn relative to the unstretched membrane length. Membranes were pre-stretched by two turns to remove slack. A silicone chamber (ibidi, 80209) with its center removed was sealed to the middle of the membrane with vacuum grease, sterilized with 70% ethanol, and coated with 10 μg/mL fibronectin in PBS for 30 min at room temperature; the coating solution was aspirated immediately before seeding. MDCKII cells were seeded in 140 μL at 8 × 10^5^ cells/mL for experiments performed 3–4 days later or at 1 × 10^6^ cells/mL for experiments performed 2–3 days later, covered with a fitted polypropylene lid, and maintained at 37 °C with medium changes every 12–24 h. Cells were labeled with 100 nM JFX_554_-HTL for 1 h and washed three times with fresh medium before imaging. For FLIM, the chamber was mounted on the microscope stage and imaged as described in **FLIM acquisition**. For each experiment, 10–20 fields of view were acquired as the pre-stretch condition. The membrane was then stretched by ∼100% engineering strain relative to the pre-stretched state and held for 10 min before 10–20 fields of view were acquired as the stretched condition. An equal volume of medium containing 2 μM latrunculin A (MilliporeSigma, 428026) was added to give a final concentration of 1 μM, and 10–20 fields of view were acquired 30 min later as the stretched + LatA condition.

### HaloTag labeling of *C. elegans*

Worms were incubated in 250 μL PCR tubes containing 20 μM JFX_554_-HTL in S-medium supplemented with excess bacterial food. Excess bacteria were confirmed by persistent turbidity of the labeling solution throughout the entire labeling period. OP50 *E. coli* was used for most experiments. For control and *plc-1* RNAi experiments, worms were labeled in the presence of HT115 bacteria carrying either the empty L4440 vector or the *plc-1* RNAi construct. After labeling, worms were recovered on NGM plates seeded with the corresponding bacterial strain. Labeling and recovery times were optimized for each sensor line: HMR-1/CeEcad worms were labeled for 8–12 h and recovered for less than 2 h; DEB-1/CeVin worms were labeled for 8–12 h and recovered for 2–6 h; and LAM-2/CeLam worms were labeled for 6–12 h and recovered for at least 2 h for spermatheca, or up to 48 h for pharynx imaging.

### Mounting *C. elegans* for FLIM imaging

Worms were mounted immediately before each imaging session. For still imaging, molten 2% (w/v) agarose was flattened on a glass slide with a second slide held parallel, using two layers of laboratory tape as spacers to set a uniform pad thickness. Five to eight worms at the indicated stage were anesthetized in a droplet of M9 buffer containing 10 mM tetramisole (MilliporeSigma, T1512), rinsed once in M9 buffer, transferred to the center of the solidified pad, and covered with a glass coverslip.

For time-lapse imaging of ovulation, worms were immobilized in a UV-curable mountant. One drop of BIO-133 (MY Polymers) was placed at the center of a coverslip. Two to four worms were anesthetized as above, rinsed once in M9 buffer, and transferred into the droplet with minimal M9 carryover. Worms were pressed gently against the coverslip with a worm pick. A UV flashlight (uvBeast V1, 385–395 nm) was positioned one glass slide thickness (∼1 mm) above the coverslip and the mountant was cured for 40–60 s, confirmed by gently probing the droplet with a worm pick. The coverslip was then mounted onto a glass slide with a 0.12 mm spacer (Grace Bio-Labs, 654002).

### Estimation of ovulation number

Worms were staged precisely at the L4 stage, labeled immediately with JFX_554_-HTL for at least 6 h at 20 °C, and recovered overnight at 16 °C for approximately 12 h. The following morning, these day-1 adults contained no more than one embryo in the uterus and no eggs were present on the plates, indicating that egg laying had not yet begun. Individual worms were singled immediately before imaging, and their plates were checked to confirm that no eggs were laid. Because no eggs had been laid, the number of embryos retained on each side of the vulva was used to infer the cumulative number of ovulations from the corresponding spermatheca. Embryos anterior and posterior to the vulva were counted separately, yielding an ovulation number for each of the two spermath-ecae. The anterior and posterior spermathecae belong to anatomically separate gonad arms and constitute distinct local mechanical structures, each with its own ovulation history and basement-membrane lifetime measurement.

Accordingly, the spermatheca was used as the unit of analysis for the correlation between ovulation number and local CeLam-WF lifetime contrast. Ovulation continued over the several hours of each imaging session, so animals imaged later had accumulated more ovulations, spanning 0 to 9 across the dataset.

### Protein expression, purification, and labeling

cpHalo and cpWHalo proteins bearing flanking ALFA tags and a C-terminal His tag were expressed from IPTG-inducible pET plasmids synthesized by Twist Bioscience. Plasmids were transformed into T7 Express *E. coli* (NEB, C2566I), and expression was induced at an OD_600_ of 0.5–0.8 with 500 μM IPTG (MilliporeSigma, I6758) for 2 h at 37 °C. Cells were harvested, lysed in B-PER (Thermo Fisher, 89821) supplemented with Halt protease inhibitor cocktail (Thermo Fisher, 78438), and clarified by centrifugation at 30,000 × g for 1 h at 4 °C. His-tagged proteins were purified by nickel-affinity chromatography (Qiagen, 30210). The resin was equilibrated in buffer containing 50 mM HEPES, pH 7.2, and 250 mM NaCl; after protein binding, the resin was washed in H250 buffer (25 mM HEPES, pH 7.5, 250 mM NaCl) with 25 mM imidazole, and proteins were eluted in H250 buffer containing 250 mM imidazole. DTT was added to the eluates at a final concentration of 1 mM. Where indicated, nickel-purified proteins were buffer-exchanged into H250 buffer and further purified using peptide-elutable ALFA-tag affinity resin (NanoTag, N1515-L). Purity was assessed by SDS-PAGE and Coomassie staining. Selected fractions were pooled, buffer-exchanged into H250 buffer, aliquoted, snap-frozen in liquid nitrogen, and stored at ™80 °C. For dye labeling, purified cpHalo and cpWHalo proteins were diluted to 2–5 μM in H250 buffer and incubated with 20 μM JFX_554_-HTL (from 10 mM stock) for 2 h on an end-to-end rotator. Excess dye was removed by buffer exchange using the Zeba spin desalting column following the manufacturer’s instructions (Thermo Fisher, 89883 or 89890).

### Photophysical characterization

Purified cpHalo and cp-WHalo proteins in H250 buffer (25 mM HEPES, pH 7.5, and 250 mM NaCl) were substoichiometrically labeled with 2.5 μM JFX_554_-HTL in the presence of excess protein for 1 h while protected from light. A 2.5 μM JFX_554_-HTL sample in H250 buffer was prepared as a dye-only control. Absorbance spectra from 200 to 800 nm were acquired using an Agilent Cary Series UV-visible spectrophotometer with H250 buffer as the reference. Molar extinction coefficients were determined from the slopes of absorbance versus concentration using the Beer-Lambert law across the undiluted sample and two successive 1:2 dilutions. Fluorescence excitation and emission spectra were acquired using an Agilent Cary Eclipse fluorescence spectrophotometer. Excitation spectra were collected from 480 to 640 nm with emission monitored at 650 nm, and emission spectra were collected from 530 to 680 nm with excitation at 520 nm; excitation and emission slit widths were 5 nm. Absolute fluorescence quantum yields were measured using a Quantaurus-QY spectrometer (Hamamatsu, C11374) with excitation wavelengths from 500 to 530 nm at 5-nm intervals and H250 buffer as the reference.

### DNA handle functionalization for single-molecule force spectroscopy

A 7.5 kb DNA handle containing a dual-biotin (Duobiotin) at one end and an ALFAtag nanobody on the other end was prepared. First, PCR amplification was performed with a biotinylated primer (Duobiotin-5’-TCATCTGAAACAGCAGCGGA-3’, IDT) and an azide functionalized primer (Azide-5’-CGCACGAAAAGCATCAGGTC-3’, IDT) using Lambda DNA as a template. After purification of the DNA handle with a PCR cleanup kit (Qiagen), the Duobiotin-7.5kb-DNA-Azide handle was mixed with DBCO-ALFA-Nb (Nan-oTag, N1505-DBCO) at a 1:30 molar ratio and incubated overnight at 16 °C under 500 rpm agitation. Following anion-exchange purification to remove free DBCO-ALFA-Nb, the DNA handle was conjugated with cpWHalo-554 or cpHalo-554 by mixing at a 2:1 molar ratio in H250B buffer (50 mM HEPES, 250 mM NaCl, pH 7.2) overnight at 4 °C in the dark to prevent photobleaching. The next day, samples were aliquoted without further purification, snap-frozen, and stored at ™80 °C until use.

### Single-molecule force spectroscopy using optical tweezers

Force-dependent unquenching experiments of cpWHalo-554 and cpHalo-554 were performed on a LU-MICKS C-Trap (optical tweezers combined with confocal microscopy) using a five-channel microfluidic flow cell, as described in detail previously^55^. Flow-cell channels were passivated sequentially with 0.1% BSA (Sigma, CAS:9048-46-8; 30 min), a ddH_2_O wash (10 min), 0.5% Pluronic F-127 (Sigma, CAS: 9003-11-6; 30 min), and a final 1X PBS wash (Thermo Fisher, AM9625; 30 min), all under automated flow cycles (1 bar for 5 min, then 0.4 bar for the remainder).

First, two 2.14 μm streptavidin beads (Spherotech, SVP-20-5) were optically trapped in a BSA-passivated fluid cell. The cpWHalo-554 or cpHalo-554 sample bearing two DNA handles was introduced to the beads under constant flow. By repeatedly bringing one streptavidin bead close to the other, a single-molecule tether was formed that placed cpWHalo or cpHalo between the two beads, connected through the two DNA handles (**Fig. 1e**). Single-molecule tethers that matched the expected extension of the 15 kb DNA handles (total length), according to the extensible worm-like chain model, were selected for correlated force and fluorescence measurements. Imaging was performed with 0.1–0.5 ms exposure per pixel and 100 nm pixel size, using green 575 nm excitation at ∼5 μW laser power. The collected kymo-graphs were processed through Python scripts based on the LUMICKS Pylake API. An oxygen scavenging imaging buffer containing 4 mM Trolox (Sigma-Aldrich, 238813), 2.5 mM protocatechuic acid (Sigma-Aldrich, 37580), and 0.2% (v/v) rPCO (∼0.01 units/mL; OYC Americas, 46852004) was used in H250 buffer. The time it took until cpWHalo-554 showed increased fluorescence intensity due to unquenching was measured as dwell time and quantified. Flow cells were cleaned between experiments with ddH_2_O, 2.5% bleach (Clorox), a water rinse, sodium thiosulfate neutralization (Millipore, CAS:7772-98-7; 10 mM), and a final water wash, then re-passivated for subsequent use.

### Estimation of force-dependent unquenching rates

Quenched-state dwell-time distributions were visualized as Kaplan–Meier survival curves. For each constant-force condition, a quenched-to-unquenched transition was treated as an observed event, whereas an observation ending before transition because of photobleaching, tether breakage, or the planned acquisition endpoint was treated as right-censored. The force-dependent unquenching rate was estimated under a single-exponential model as *k*_QU_ = *n*_QU_/*T* _exp_, where *n*_QU_ is the number of observed transitions and *T* _exp_ = ∑_*i*_*t*_*i*_ is the total quenched-state observation time across both event and right-censored intervals. The corresponding mean quenched-state dwell time was calculated as τ_Q_ = *T* _exp_/*n*_QU_ = 1/*k*_QU_. Two-sided 95% exact Poisson confidence intervals were calculated using Garwood limits: *k*_low_ = χ^2^_0.025_(2*n*_QU_)/(2*T* _exp_) and *k*_high_ = χ^2^_0.975_(2(*n*_QU_+1))/(2*T* _exp_). When no transition was observed, no point estimate was assigned; instead, the one-sided 95% upper confidence bound was calculated as *k*_QU,upper_ = ™ln(0.05)/*T* _exp_. Calculations were performed in Python using a custom script with NumPy, pandas and SciPy.

### FLIM acquisition

Time-correlated single-photon-counting fluorescence lifetime imaging microscopy (TCSPC-FLIM) was performed using a Leica STELLARIS microscope built on a DMi8 inverted stand and controlled with Leica Application Suite X (LAS X v4.5.0.25531). Images were acquired using a 40X 1.25 NA glycerol-immersion objective (Leica, 11506422). Fluorescence was detected using a fast-response HyD X2 or HyD X4 detector, depending on the experimental configuration.

Samples were excited with an 80-MHz pulsed white-light laser. Unless otherwise indicated, excitation was performed at 561 nm, and emitted fluorescence was collected from 565 to 650 nm. Excitation power, scan speed and signal accumulation were adjusted for each sample to obtain sufficient photon counts while minimizing photobleaching and avoiding detector saturation and photon pile-up. Acquisition settings were chosen to provide at least 100 detected photons per pixel at the structures of interest. For exceptionally dim samples, a minimum threshold of 50 photons per pixel was accepted. Detector identity, spectral window and acquisition settings were held constant within each experiment and matched between experimental groups.

For live imaging of cultured cells and salivary gland explants, samples were maintained in a Tokai Hit stage-top incubation system at 37 °C and 5% CO_2_. Humidity was maintained passively by filling the incubator’s water reservoir before imaging. Environmental conditions were maintained throughout image acquisition. *C. elegans* were imaged at ambient room temperature (∼22 °C) without environmental control.

### Quantitative fluorescence lifetime analysis

Fluorescence decays were analyzed in Leica LAS X (v4.8.1.29271) by reconvolution fitting to a two- or three-exponential decay model, as appropriate for the fluorophore and experiment. The fit returned the component lifetimes (τ_i_), amplitudes (α_i_) and amplitude- and intensity-weighted mean lifetimes. The amplitude-weighted mean lifetime, which is proportional to the fluorescence quantum yield, was calculated as: τ_amp_ = (∑_*i*_α_*i*_τ_*i*_)/(∑_*i*_α_*i*_). The intensity-weighted mean lifetime, which weights each component by its contribution to the total emission, was calculated as: τ_int_ = (∑_*i*_α_*i*_τ_*i*_^2^)/(∑_*i*_α_*i*_τ_*i*_).

For spatial and region-of-interest (ROI) analyses, time-correlated single-photon-counting data were exported from LAS X as PTU files and processed using custom Python scripts. Cells, focal adhesions or epithelial junctions were segmented from fluorescence-intensity images using Cellpose-SAM 4.1.1^56,57^. Depending on the assay, segmentation used either the base Cellpose-SAM model or models fine-tuned on representative images using the Cellpose graphical user interface^58^. Separate fine-tuned models were used when perturbations substantially altered object morphology. Segmentation produced integer label images in which each cell or subcellular structure was assigned a unique ROI.

The instrument response function (IRF) was recovered from a Rhodamine B reference (2 μM in ethanol) acquired at 37 °C or room temperature with detector and spectral settings matched to the corresponding experiment. The reference was fitted in LAS X against the instrument IRF to obtain its measured mono-exponential lifetime; dataset-specific fitted values ranged from approximately 2.088 to 2.196 ns. The experimental IRF was then estimated from the reference decay by Gaussian-damped Richardson-Lucy deconvolution against this fitted lifetime. Where necessary, an additional acquisition-specific temporal shift was determined by minimizing the reconvolution residual against experimental decays.

For each image, the component lifetimes obtained from the corresponding LAS X fit were held fixed. The amplitudes of the IRF-convolved exponential components and a constant-background term were estimated by non-negative least squares (NNLS; scipy.optimize). For quantitative ROI measurements, the raw photon decays of all qualifying pixels within an ROI were summed and fitted once, rather than averaging independently fitted pixel lifetimes. This pooled-decay approach increased the photon budget and reduced the downward bias caused by non-negativity constraints in low-photon pixels. Per-pixel lifetime maps were used for visualization and spatial analyses, whereas pooled-decay NNLS estimates were used as the principal ROI-level quantitative measurements. Photon-count, ROI-size, and fit-quality thresholds were specified for each assay, ranging from a few hundred to several thousand photons per ROI. Fields of view were excluded for low photon counts, poor fit quality, or imaging artifacts such as aggregates. Data were pooled across independent imaging sessions.

### Super-resolution microscopy and image processing

Super-resolution images of salivary gland samples were acquired on a home-built line-rescan confocal system^59,60^, to be fully described in a forthcoming manuscript. We used a 60X 1.2 NA water immersion objective (Nikon, CFI Plan Apo VC 60XC WI) to excite the sample with 405 nm, 488 nm, 561 nm, and 637 nm laser lines (Coherent, OBIS CellX). Four bandpass emission filters (Semrock FF02-447/60, FF03-525/50, FF01-600/52, and FF01-680/42) were used to block the excitation lasers and select the corresponding spectral bands. The rescan confocal images were collected by a scientific complementary metal-oxide-semiconductor (sCMOS) camera (Hamamatsu, ORCA-Fusion BT). Images were acquired with a pixel size of 53.5 nm, an axial step size of 200 nm, and a frame time of 40 ms per channel. All images were deconvolved using 30 iterations of the Richardson-Lucy algorithm^61,62^ in MATLAB (code available at https://github.com/eguomin/regDeconProject).

### Brood-size and embryonic-viability assays

Brood size was quantified for each strain at 20 °C. For each strain, more than 20 L4 hermaphrodites were picked onto a single plate on Day 0. On Day 1 (24 h later), 10–15 adult worms were singled onto individual plates for each experiment, and results were pooled across experiments. On Day 2 (exactly 24 h after singling), the mother worm was removed from each plate. Plates where the mother worm was dead, missing, or trapped at the plate margin were excluded from analysis. On Day 3 (48 h after singling), hatched and unhatched embryos on each plate were counted. A minimum of 12 plates per strain were scored.

### Statistics

The independent experimental unit and definition of n are reported in each figure legend. For single-molecule measurements, each independently formed molecular tether was one experimental unit. For cultured-cell imaging, inference was performed at the cell or field-of-view (FOV) level, as specified for each assay; pixels, focal adhesions, junction segments, radial bins, and repeated time points within the same cell or FOV were nested measurements and were not treated as independent biological replicates. Experiments were replicated across separate culture preparations, dishes or transduction batches. Each cardiomyocyte time series was one experimental unit. One FOV was acquired per cardiomyocyte, so FOV and cell counts are equivalent for these experiments. For *C. elegans* experiments, the animal was the unit of analysis except for the ovulation-lifetime correlation, where each spermatheca was treated as a separate unit of analysis. Correlation between local lifetime contrast at the spermathecal basement membrane and cumulative ovulation number was quantified by Pearson r, with a 95% confidence interval estimated by whole-worm bootstrap resampling. For mouse experiments, each independently derived embryo, animal or tissue preparation was one biological replicate, as specified in the corresponding legend. Sample sizes were selected on the basis of prior experience with the assays, the availability of biological material and consistency with related studies; no formal prospective power calculations were performed. Automated analysis parameters were applied uniformly within each experiment.

Unless otherwise stated, statistical tests were performed on independent cell-, FOV-, animal- or molecule-level summaries and were two-sided. Two groups were compared using Mann-Whitney U tests, paired t-tests for within-animal comparisons between two regions of the same worm, or Welch’s t-tests for unequal variances; the test used for each comparison is stated in the corresponding figure legend. Family-wise error was controlled using the Holm-Bonferroni procedure across the related comparisons identified in each analysis. Statistical significance was defined as p < 0.05 after adjustment where applicable.

Time-dependent lifetime responses were reduced to pre-specified trajectory-level summaries, including the trapezoidal area under the baseline-subtracted lifetime curve and the median or endpoint response within the indicated time window. Quenched-state dwell-time distributions were visualized using Kaplan-Meier survival estimates. Force-dependent unquenching rates *k*_QU_, confidence intervals and zero-event upper bounds were calculated as described under **Estimation of force-dependent unquenching rates**.

For cardiomyocyte analyses, the fraction of cells exhibiting significant lifetime-beating coupling was compared between constructs using a two-sided Fisher’s exact test. Lifetime-beating coupling within each cell was quantified as the maximum normalized cross-correlation within ± one-half beating cycle after detrending and band-pass filtering. Significance was evaluated using 1,000 phase-randomized lifetime surrogates, which preserved the lifetime trace’s Fourier amplitudes and hence its power spectrum while randomizing phase. The complete filtering and lag-optimization procedure was repeated for every surrogate. The one-sided Monte Carlo p value was calculated as: *p* = (1 + #{*r*_*surrogate*_ ≥ *r*_*observed*_})/(*N*_*surr*_ + 1), where *N*_*surr*_ = 1000. A fixed random-number seed of 12345 was used.

## Code availability

Analysis was performed in Python 3.11 using NumPy, SciPy, pandas, scikit-image, tifffile, ptufile, lmfit, statsmodels, Matplotlib and Cellpose 4.1.1; exact versions are specified in the repository environment file. Fluorescence-decay reconvolution fits were performed in Leica LAS X (v4.8.1.29271). Custom analysis code and reproduction commands are available at https://github.com/shaohewanglab/WHaloForce and archived at Zenodo (https://doi.org/10.5281/zenodo.21895937).

## Data availability

Source data underlying all figures are provided as CSV files in the code repository. Raw FLIM data (PTU files) and processed intermediates will be deposited at Figshare and released upon publication (DOI 10.25378/janelia.33213720).

## Materials availability

Plasmids are available on Addgene (IDs #261833–#261846). The *C. elegans* strains listed in **Table S1** will be deposited at CGC. The Vcl-WF mouse line will be available through corresponding-author request.

## Video Legends

**Video S1**. Time-lapse video showing 2 h of spreading in NIH-3T3 cells expressing Vin-WF under DMSO vehicle control conditions (left) or after treatment with the ROCK inhibitor Y-27632 (5 μM; right). Vin-WF was labeled with JFX_554_-HTL. Color represents intensity-weighted mean fluorescence lifetime (τ_int_). The sequence plays twice: full-field lifetime images, then only the focal-adhesion pixels used for quantification. The plot below tracks the rolling intensity-weighted mean lifetime within focal adhesions over time for each condition.

**Video S2**. Time-lapse video of a human iPSC-derived cardiomyocyte expressing Vin-WF labeled with JFX_554_-HTL. Raw lifetime is shown on the left in real time. Phase-averaged lifetime of 39 complete beat cycles, masked to the pixels with the top 25% brightness of phase-averaged photon counts, is shown on the right. Color represents intensity-weighted mean fluorescence lifetime (τ_int_). The bottom plot shows the phase-binned contraction signal and intensity-weighted mean fluorescence lifetime (± SEM) across one averaged cycle. The vertical line in the bottom plot tracks the current phase within the beating cycle.

**Video S3**. Time-lapse video of a human iPSC-derived cardiomyocyte expressing Vin-cpH labeled with JFX_554_-HTL. Raw lifetime is shown on the left in real time. Phase-averaged lifetime of 38 complete beat cycles, masked to the pixels with the top 25% brightness of phase-averaged photon counts, is shown on the right. Color represents intensity-weighted mean fluorescence lifetime (τ_int_). The bottom plot shows the phase-binned contraction signal and intensity-weighted mean fluorescence lifetime (± SEM) across one averaged cycle. The vertical line in the bottom plot tracks the current phase within the beating cycle.

**Video S4**. Time-lapse FLIM imaging of an early ovulation event (second ovulation) in a *C. elegans* hermaphrodite (CeLam-WF). CeLam-WF was labeled with JFX_554_-HTL. Top: intensity-weighted mean fluorescence lifetime images, with brightness scaled by photon count. Middle: corresponding raw photon-count images; the red line marks the traced bottom edge of the germline/spermatheca/uterus membrane sampled to build the kymograph below. Bottom: kymograph of τ_int_ along the traced membrane over time (top = t = 0), revealed progressively as the movie plays. Anatomical landmarks (Intestine, Germline, Spt = spermatheca, Uterus) are labeled on the first frame. 150 frames over 19.5 min total (∼7.8 s/frame); movie plays at 12 fps.

## Extended Data Figures

Extended Data Figures 1–7 are provided below. Supplementary Methods, Supplementary Tables and Supplementary Videos are available separately.

**Extended Data Figure 1.**
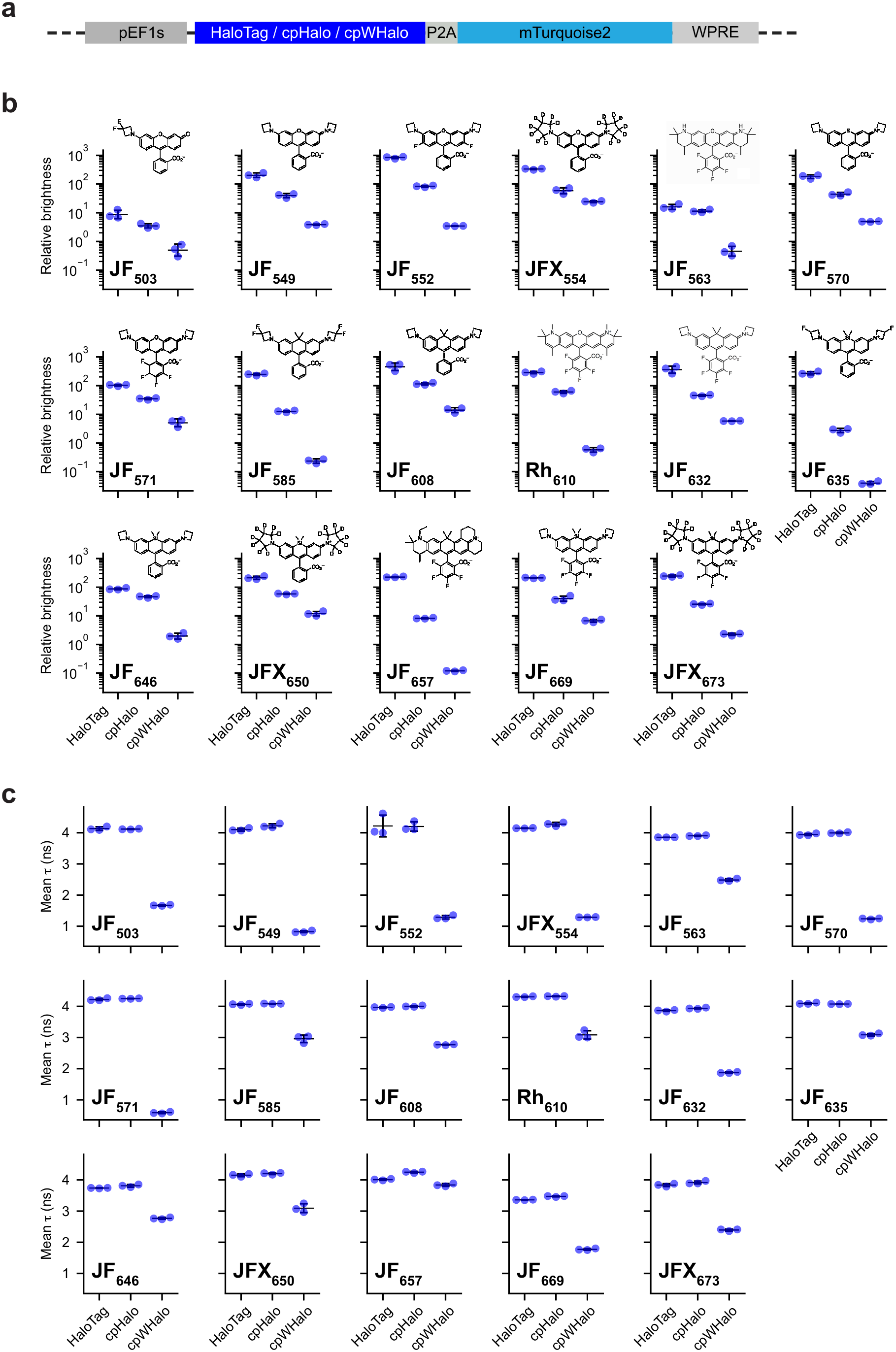
Brightness and fluorescence lifetime changes of cpHalo and cpWHalo relative to HaloTag in HEK293T cells. **(a)** Schematic of the expression cassettes for HaloTag, cpHalo, and cpWHalo. A co-translational mTurquoise2 reporter was included to normalize for expression level. **(b,c)** Per-field-of-view (FOV) relative brightness **(b)** and intensity-weighted mean fluorescence lifetime **(c)** for HaloTag, cpHalo, and cpWHalo labeled with 17 different dye-HaloTag ligands (HTLs). Free dye structures are shown in each plot in **(b)** for reference. Relative brightness was defined as total photon counts in the dye channel normalized by excitation laser power, effective acquisition time, and mTurquoise2 photon counts, which were used as a proxy for expression level. Each point represents one FOV; bars indicate mean ± SD. **Extended Data Fig. 2 a**

**Extended Data Figure 2.**
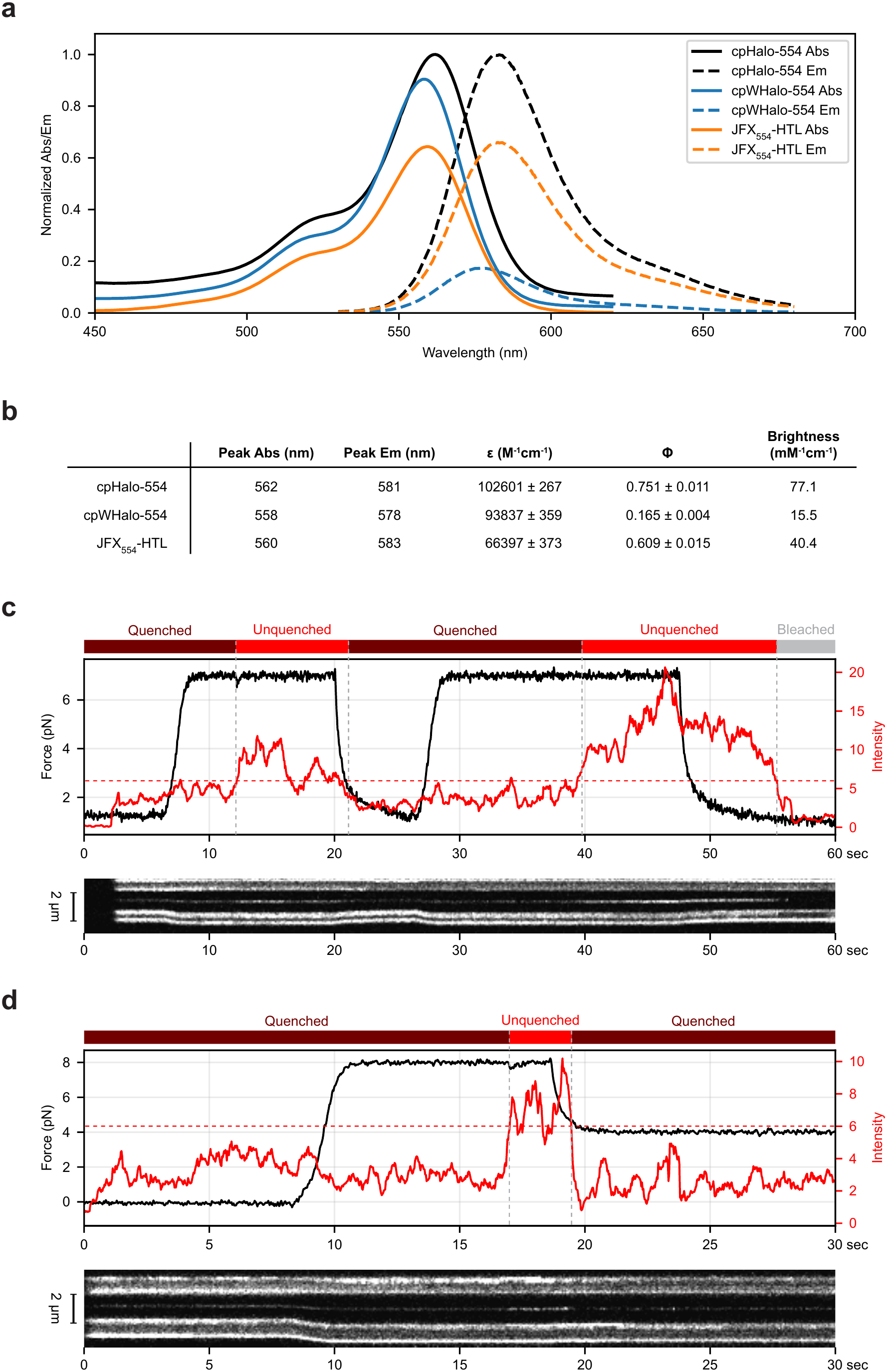
Photophysical characterization and single-molecule examples of re-quenching after force reduction. **(a)**Normalized absorption and fluorescence emission spectra of cpHalo-554, cpWHalo-554, and JFX_554_-HTL. **(b)**Table summarizing peak absorption, peak emission, molar extinction coefficient (ε), quantum yield (Φ), and molecular brightness (ε × Φ / 1,000) for cpHalo-554, cpWHalo-554, and JFX_554_-HTL. **(c,d)** Smoothed force and fluorescence intensity traces (top) and corresponding kymographs (bottom) for cpWHalo-554. Dashed red lines indicate the intensity threshold used to classify quenched and unquenched states. Colored bars above each trace indicate sensor state: quenched, unquenched, or photobleached.

**Extended Data Figure 3.**
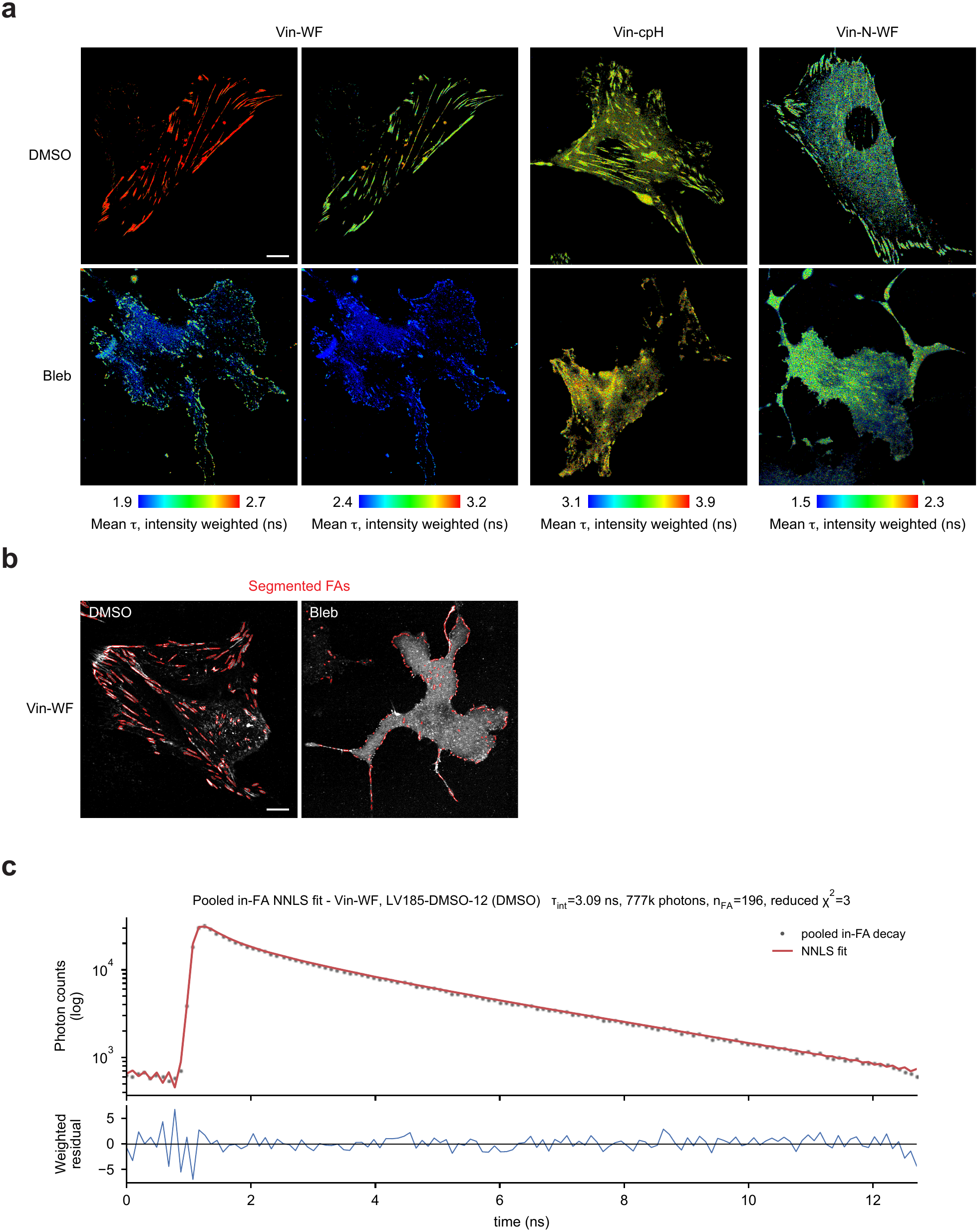
Representative FLIM images, focal-adhesion segmentation, and lifetime fitting for control and blebbistatin-treated NIH-3T3 cells. **(a)**Representative FLIM images of control and blebbistatin-treated NIH-3T3 cells expressing Vin-WF, Vin-cpH, or Vin-N-WF. Color represents intensity-weighted mean fluorescence lifetime (τ_int_). Vin-WF images are shown with two color scales. **(b)**Representative focal-adhesion (FA) segmentation masks in control and blebbistatin-treated cells, generated using a custom-tuned Cellpose-SAM model. **(c)**Representative lifetime fit and residuals using pooled photon-decay data from all in-FA pixels. The fit shown is a non-negative least-squares (NNLS) fit using an experimentally determined instrument response function (IRF) and fixed τ_1_, τ_2_, and τ_3_ values obtained from three-exponential reconvolution fits in Leica LAS X software (χ^2^ = 0.6–1.6). The higher χ^2^ value in the NNLS fit primarily reflects mismatch between the experimentally determined IRF extracted from a reference image and the image-specific IRF automatically extracted and used by LAS X during fitting. All cells were labeled with JFX_554_-HTL. Scale bars, 10 μm.

**Extended Data Figure 4.**
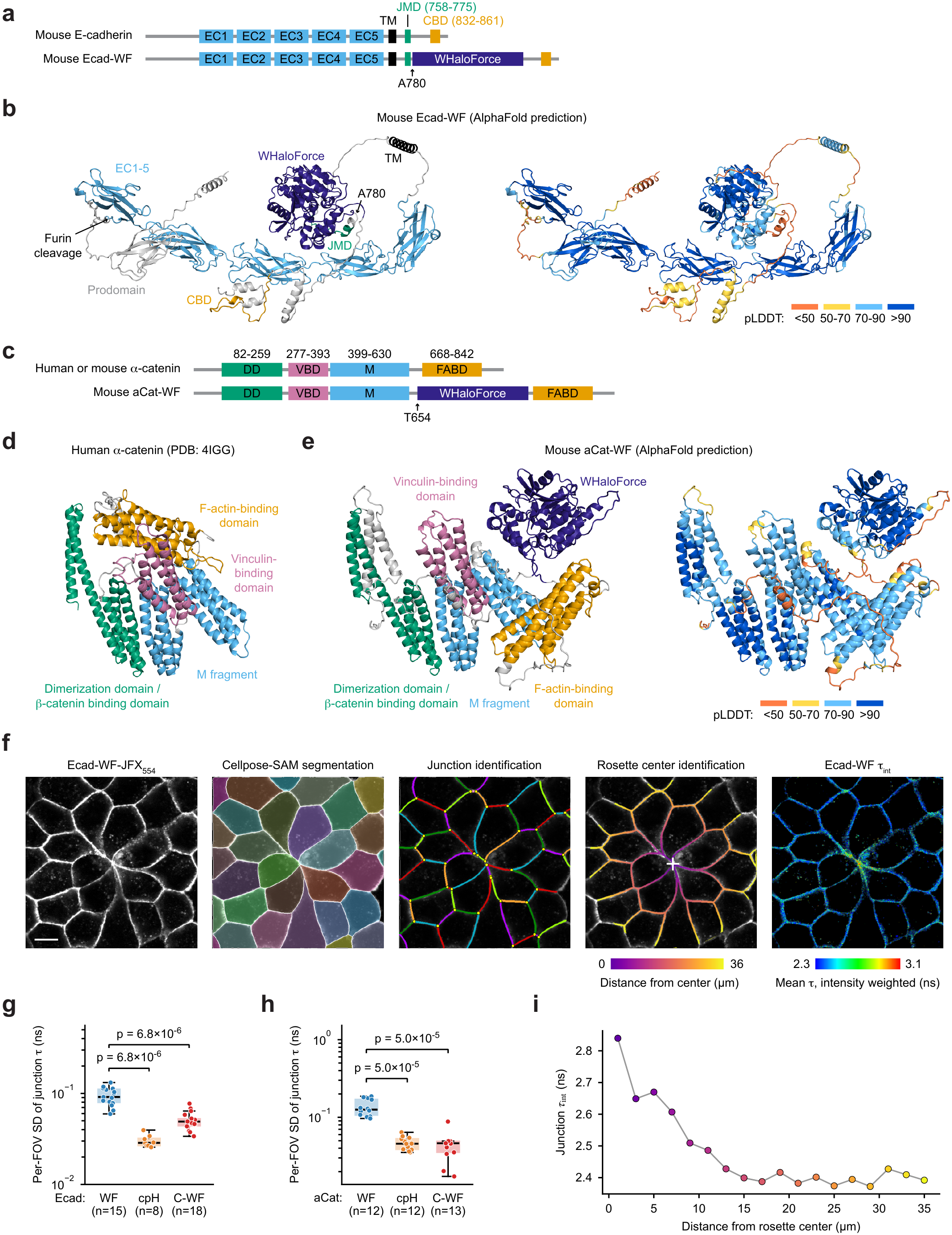
Design of E-cadherin and α-catenin sensors and adherens-junction quantification methods. **(a)**Schematics of mouse E-cadherin and Ecad-WF. EC1-5, extracellular cadherin domains 1-5, which mediate homophilic interactions between cells; TM, transmembrane domain; JMD, juxtamembrane domain that binds p120-catenin; CBD, catenin-binding domain that binds β-catenin. **(b)**AlphaFold-predicted structures of mouse Ecad-WF colored by domain (left) or pLDDT score (right). **(c)**Schematics of α-catenin (human or mouse; same amino acid numbering) and mouse aCat-WF. DD, dimerization domain; VBD, vinculin-binding domain; M, adhesion modulation domain; FABD, F-actin-binding domain. **(d)** Crystal structure of human α-catenin (PDB: 4IGG) colored by domain. **(e)** AlphaFold-predicted structures of mouse aCat-WF colored by domain (left) or pLDDT score (right). **(f)** Representative images showing Ecad-WF-JFX_554_ intensity, cell segmentation by Cellpose-SAM, junction identification, rosette-center identification, and intensity-weighted mean fluorescence lifetime. Only junctions ≥2 μm in length and containing ≥5,000 photons were included in the per-junction lifetime analysis. Cells were labeled with JFX_554_-HTL. Scale bar, 10 μm. **(g,h)** Box-and-scatter plots of per-field-of-view (FOV) standard deviation (SD) of per-junction lifetime (τ) for E-cadherin **(g)** and α-catenin **(i)** sensors and controls. Statistics: Mann-Whitney U tests with Holm-Bonferroni correction; reported p values are Holm-adjusted.**(j)** Radial profile of junction lifetime (τ) from the rosette center for the image shown in **(f)**. Each point represents the intensity-weighted mean lifetime of junctional pixels within a 2-μm-thick radial ring. Point color indicates distance from the rosette center, matching the color coding in **(f)**.

**Extended Data Figure 5.**
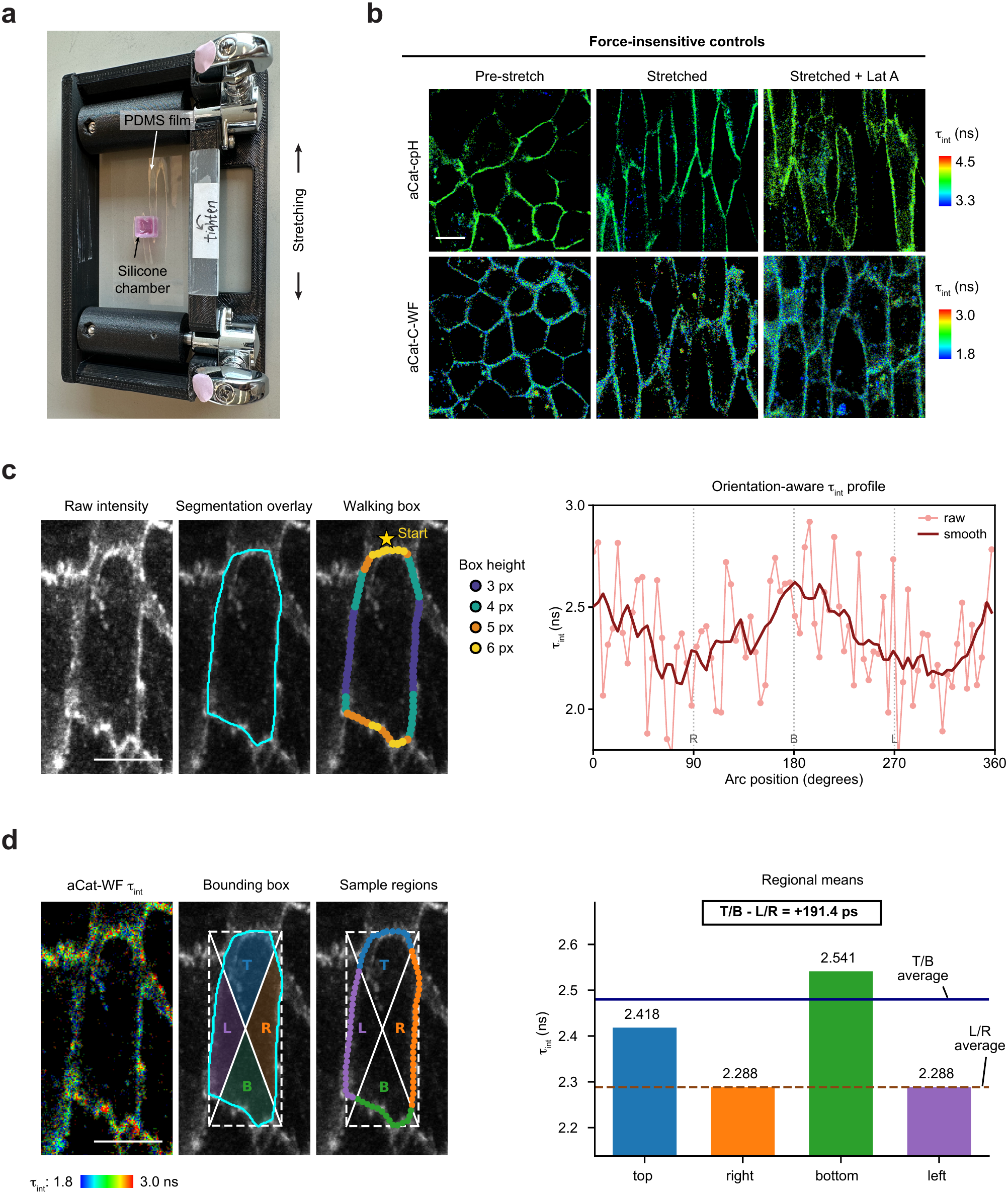
Representative FLIM images and quantification methods for the stretch experiment. **(a)** Photograph of the custom 3D-printed manual stretching device. **(b)** FLIM images of MDCKII cells expressing the aCat-cpH or aCat-C-WF controls before stretch (left), after stretch (middle), and after 1 μM latrunculin A (LatA) treatment (right). Color represents intensity-weighted mean fluorescence lifetime (τ_int_). **(c)** Representative stretched MDCKII cell expressing aCat-WF, with overlays showing the segmented cell boundary and walking boxes used to sample junctional pixels around the cell periphery (left), and corresponding plot of intensity-weighted mean fluorescence lifetime (τ_int_) along the cell periphery for the same cell (right). The walking box was advanced around the segmented cell boundary from the reference starting position; box height was adjusted according to box position to maintain consistent sampling of the junctional region along the cell perimeter. **(d)** Representative stretched MDCKII cell expressing aCat-WF, with overlays showing the segmented cell boundary, bounding box, and diagonal lines used to divide the cell into top (T), bottom (B), left (L), and right (R) regions (left), and corresponding plot of intensity-weighted mean fluorescence lifetime (τ_int_) in the four regions for the same cell (right). These regions were used to calculate junctional lifetime anisotropy, Δτ, defined as the difference between the mean lifetime of pooled junction pixels in the top and bottom regions and the mean lifetime of pooled junction pixels in the left and right regions. All cells were labeled with JFX_554_-HTL. Scale bars, 10 μm.

**Extended Data Figure 6.**
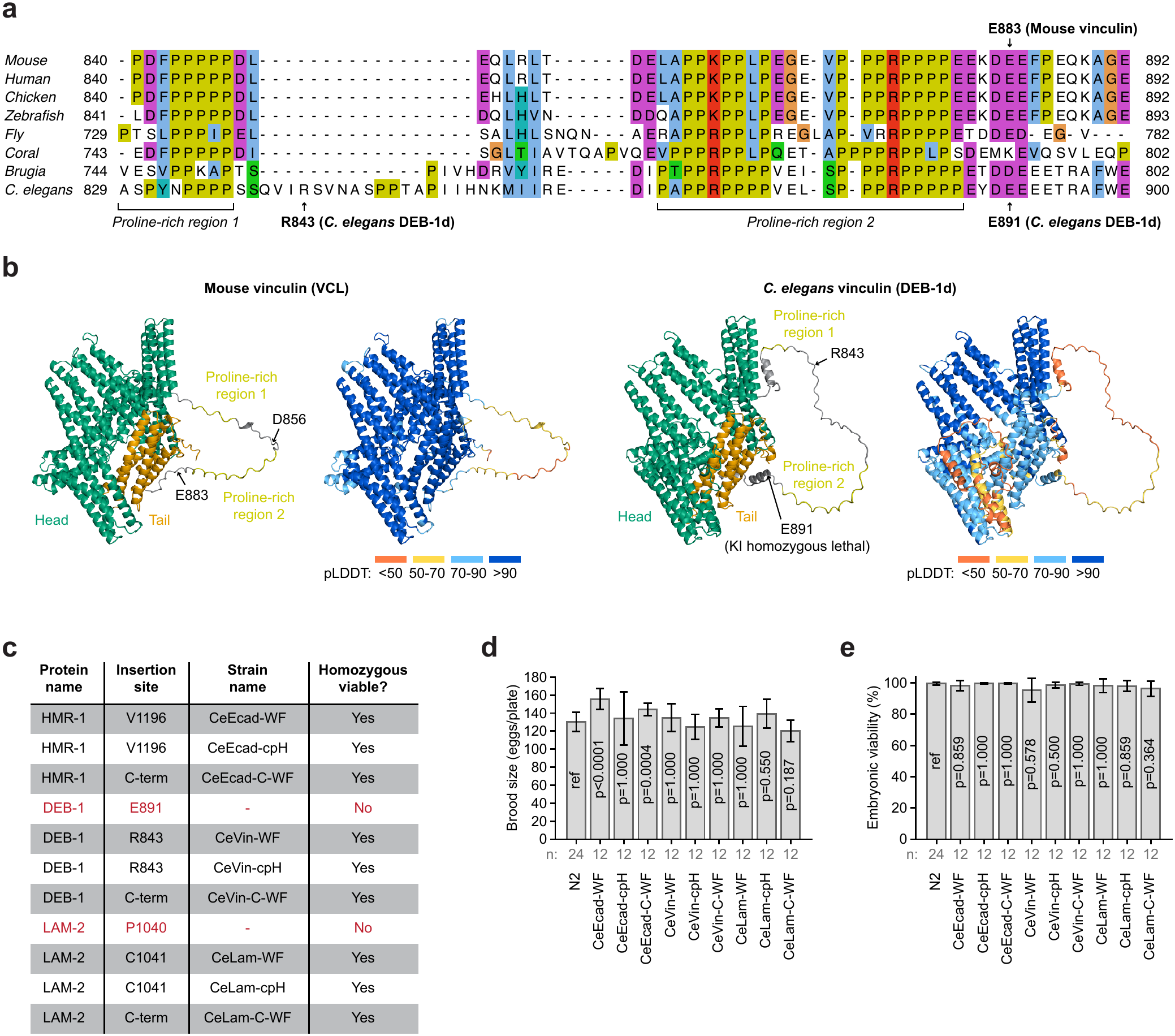
Design of the *C. elegans* vinculin sensor and phenotypic characterization of *C. elegans* knock-ins. **(a)** Protein sequence alignment of the vinculin linker region from the indicated species. Multiple sequence alignment was performed in Jalview using the T-Coffee method. Colors indicate conserved residue classes according to the ClustalX coloring scheme in Jalview. NCBI accession numbers for the sequences used are NP_033528.3 (mouse), NP_003364.1 (human), XP_015143689.1 (chicken), NP_001122153.1 (zebrafish), CAA65421.1 (fly), XP_022799230.1 (coral), XP_001899040.1 (*Brugia malayi*), and NP_001293743.1 (*C. elegans*). **(b)** AlphaFold-predicted structures of mouse vinculin (left) and *C. elegans* vinculin/DEB-1d (right), colored by domain or pLDDT score as indicated. **(c)** Table summarizing viability outcomes for *C. elegans* knock-in lines at the indicated insertion sites in E-cadherin/HMR-1, vinculin/DEB-1, and laminin γ1/LAM-2. HMR-1 amino acid numbering corresponds to isoform a. DEB-1 amino acid numbering corresponds to isoform d. **(d)** Brood size from 24 to 48 h post-L4 for the indicated *C. elegans* strains. **(e)** Embryonic viability of embryos laid from 24 to 48 h post-L4 by the indicated *C. elegans* strains. Statistical significance for panels d and e was assessed by two-sided Welch’s t-test comparing each strain to N2, with Holm-Bonferroni correction for multiple comparisons. Bars in panels d and e indicate mean ± SD.

**Extended Data Figure 7.**
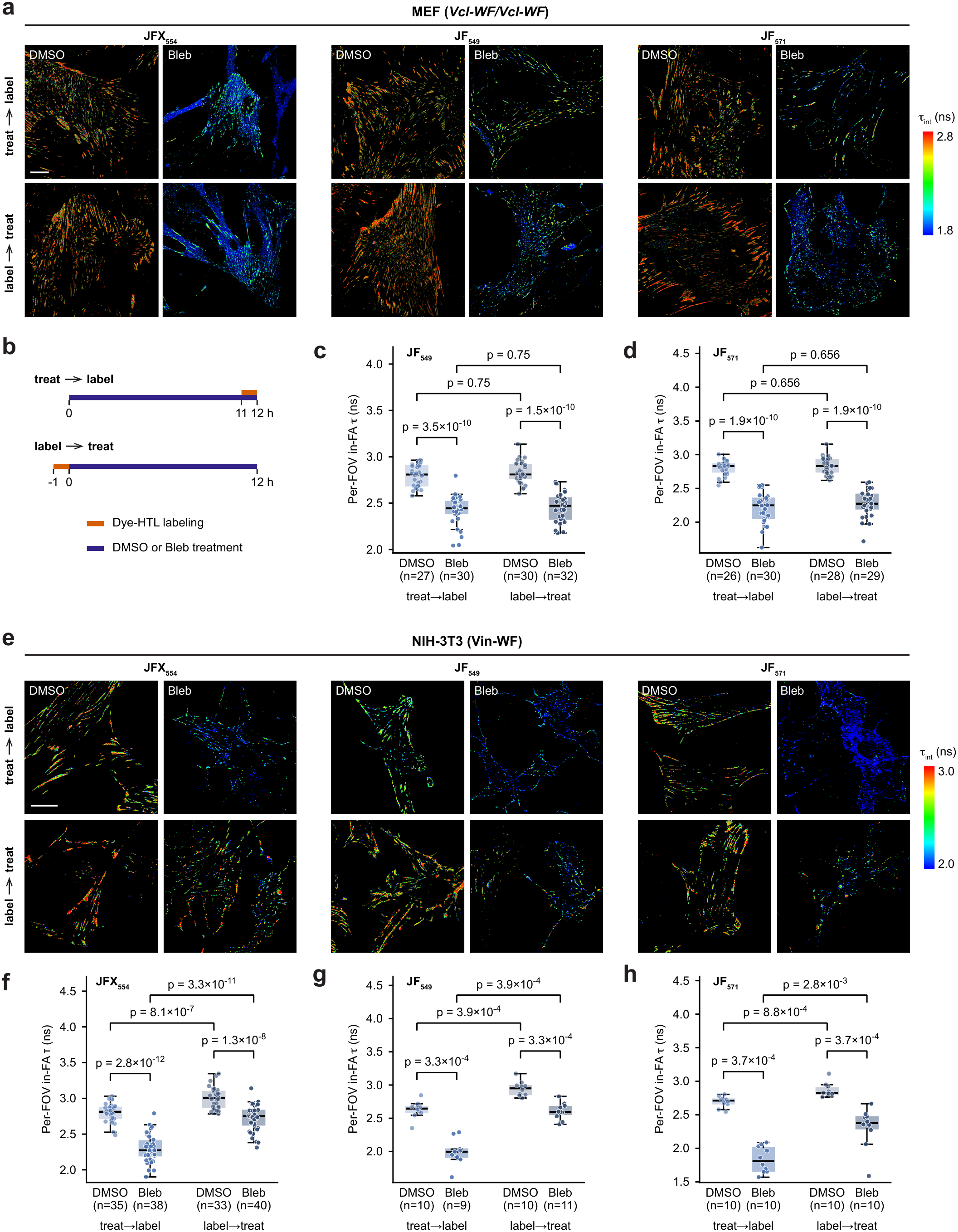
Staged labeling of endogenous Vcl-WF in MEFs and of ectopically expressed Vin-WF in NIH-3T3 cells with three dye-HTLs. **(a)** FLIM images of *Vcl-WF/Vcl-WF* MEFs labeled with JFX_554_-, JF_549_-, or JF_571_-HTL under the indicated conditions. Color represents intensity-weighted mean fluorescence lifetime (τ_int_). **(b)** Staged-labeling timeline, as in Figure 5d. **(c,d)** Box-and-scatter plots of per-FOV mean in-FA lifetime of MEFs labeled with JF_549_-HTL (c) or JF_571_-HTL (d). Data were pooled from two independent experiments. **(e)** FLIM images of NIH-3T3 cells expressing Vin-WF, labeled with JFX_554_-, JF_549_-, or JF_571_-HTL under the indicated conditions. Color represents intensity-weighted mean fluorescence lifetime (τ_int_). **(f–h)** Box-and-scatter plots of per-FOV mean in-FA lifetime of NIH-3T3 cells expressing Vin-WF and labeled with JFX_554_-HTL (f; pooled from three independent experiments), JF_549_-HTL (g; one experiment), or JF_571_-HTL (h; one experiment). Each point represents one FOV; different FOVs contain different cells. Statistics: Mann-Whitney U tests with Holm-Bonferroni correction; reported p values are Holm-adjusted. Scale bars, 20 μm.

