## Supplementary Information for "WHaloForce enables chemigenetic imaging of molecular tension in living cells and animals"

Di Wu, E. Angelo Morales, Joon Lee, Sushil Pangeni, Helen Farrants, Katherine A. Hutchings, Robert J. Barndt, Qi Yu, Xuesong Li, Hari Shroff, Haodi Wu, Alison G. Tebo, Luke D. Lavis, Eric R. Schreiter, Taekjip Ha, Shaohe Wang

**Supplementary Note 1: cpHalo and cpWHalo protein sequences**

**Sequence of cpHalo used in this work:**

VLTEVEMDHYREPFLNPVDREPLWRFPNELPIAGEPANIVALVEEYMDWLHQSPVPKLLFWGTP
GVLIPPAEAARLAKSLPNCKAVDIGPGLNLLQEDNPDLIGSEIARWLSTLEISGGGTGGSGGTGG
SGGTGGSAEIGTGFPFDPHYVEVLGERMHYVDVGPRDGTPVLFLHGNPTSSYVWRNIIPHVAP
THRCIAPDLIGMGKSDKPDLGYFFDDHVRFMDAFIEALGLEEVVLVIHDWGSALGFHWAKRNPE
RVKGIAFMEFIRPIPTWDEWPEFARETFQAFRTTDVGRKLIIDQNVFIEGTLPMGVVR

**Sequence of cpWHalo/WHaloForce used in this work:**

VLTEVEMDHYREPFLNPVDREPLWRFPNELPIAGEPANIVALVEEYMDWLHQSPVPKLLFWGTP
GVLIPPAEAARLAKSLPNCKAVDIGPGLNLLQEDNPDLIGSEIARWLSTLEISGGGTGGSGGTGG
SGGTGGSAEIGTGFPFDPHYVEVLGERMHYVDVGPRDGTPVLFLHGNPTSSYVWRNIIPHVAP
THRCIAPDLIGMGKSDKPDLGYFFDDHVRFMDAFIEALGLEEVVLVIHDWGSALGFHWAKRNPE
RVKGIAFMEFIRPIPTWDEWPEFARETFQAFRTTDVGRKLIIDQNVFIE**W**TLPMGV**W**R

**Notes:**

GGTGGSGGTGGSGGTGGS: circular permutation linker

**V**, **W**, **W**: P180V, G171W, and V178W, in order of appearance. All carried over from prior HaloTag optimization work.

**W** (G171W): quenching tryptophan.

**Table S1. *C. elegans* strains used or generated in this study.**

| Strain ID | Genotype | Strain name | Source |
|---|---|---|---|
| N2 | *wild type (ancestral)* | — | CGC |
| DDD1 | *lam-2(bbb1[LAM-2(C1041)::WHaloForce]) X* | CeLam-WF | This study |
| DDD4 | *lam-2(bbb4[LAM-2(C1041)::cpHalo]) X* | CeLam-cpH | This study |
| DDD47 | *lam-2(bbb28[LAM-2::WHaloForce]) X* | CeLam-C-WF | This study |
| DDD37 | *lam-2(bbb26[LAM-2(P1040)::WHaloForce])/+ X* † | — | This study |
| DDD25 | *deb-1(bbb20[DEB-1(R843)::WHaloForce]) IV* | CeVin-WF | This study |
| DDD29 | *deb-1(bbb24[DEB-1(R843)::cpHalo]) IV* | CeVin-cpH | This study |
| DDD50 | *deb-1(bbb31[DEB-1::WHaloForce]) IV* | CeVin-C-WF | This study |
| DDD5 | *deb-1(bbb5[DEB-1(E891)::WHaloForce])/+ IV* † | — | This study |
| DDD9 | *hmr-1(bbb9[HMR-1(V1196)::WHaloForce]) I* | CeEcad-WF | This study |
| DDD18 | *hmr-1(bbb17[HMR-1(V1196)::cpHalo]) I* | CeEcad-cpH | This study |
| DDD58 | *hmr-1(bbb29[HMR-1::WHaloForce]) I* | CeEcad-C-WF | This study |
| DZ325 | *ezIs2 III; him-8(e1489) IV* | — | CGC |
| DDD39 | *lam-2(bbb1[LAM-2(C1041)::WHaloForce]) X; ezIs2 III* | — | This study |
| DDD40 | *lam-2(bbb4[LAM-2(C1041)::cpHalo]) X; ezIs2 III* | — | This study |
| DDD57 | *lam-2(bbb28[LAM-2::WHaloForce]) X; ezIs2 III* | — | This study |
| CF1903 | *glp-1(e2144) III* | — | CGC |
| DDD52 | *lam-2(bbb1[LAM-2(C1041)::WHaloForce]) X; glp-1(e2144) III* | — | This study |
| DDD53 | *lam-2(bbb4[LAM-2(C1041)::cpHalo]) X; glp-1(e2144) III* | — | This study |
| DDD56 | *lam-2(bbb28[LAM-2::WHaloForce]) X; glp-1(e2144) III* | — | This study |

†The *bbb26* and *bbb5* alleles are homozygous lethal. Each line was cryopreserved as a mixed population of wild-type and heterozygous animals; neither was used experimentally, and both are listed only to document their construction.

**Table S2. Genotyping primers for *C. elegans* and mouse knock-ins.**

| Strain | PCR assay | Primer | ID | Sequence (5′→3′) | Amplicon size |
|---|---|---|---|---|---|
| **CeLam-WF** | Left junction PCR | F | oS313 | ACTTCTGGACTTGGATGCCA | 772 bp |
| **CeLam-cpH** | | R | oS314 | AGTGTGGGTCGAATGGGAAT | |
| | Right junction PCR | F | oS315 | AGTTCATCCGTCCAATCCCA | 700 bp |
| | | R | oS316 | CGTGTTCCAGAAGCCAACTC | |
| | Three-primer PCR | F | oS309 | TGTCAACCATGCGATTGTGA | 313 bp (WT); 613 bp (KI) |
| | | R | oS314 | AGTGTGGGTCGAATGGGAAT | |
| | | R | oS310 | CTTCTGAAGCAGCACGAGAC | |
| **CeLam-C-WF** | Left junction PCR | F | oD29 | GATACCATGGAAGAAGTGAAC | 812 bp |
| | | R | oS314 | AGTGTGGGTCGAATGGGAAT | |
| | Right junction PCR | F | oS315 | AGTTCATCCGTCCAATCCCA | 524 bp |
| | | R | oD30 | GAATGGCTCAATTGGGTTG | |
| | Three-primer PCR | F | oD29 | GATACCATGGAAGAAGTGAAC | 688 bp (WT); 812 bp (KI) |
| | | R | oS314 | AGTGTGGGTCGAATGGGAAT | |
| | | R | oD30 | GAATGGCTCAATTGGGTTG | |
| **CeVin-WF** | Left junction PCR | F | oS429 | GAAGGTTGAGGACTGTGTTTCG | 805 bp |
| **CeVin-cpH** | | R | oS314 | AGTGTGGGTCGAATGGGAAT | |
| | Right junction PCR | F | oS315 | AGTTCATCCGTCCAATCCCA | 499 bp |
| | | R | oS430 | CCTTCACCTCTAACCAACTG | |
| | Three-primer PCR | F | oS465 | CCCTCCAAGATCGCCTCTCTC | 237 bp (WT); 617 bp (KI) |
| | | R | oS466 | CGTGGTGGAGCTGGAATATC | |
| | | R | oS314 | AGTGTGGGTCGAATGGGAAT | |
| **CeVin-C-WF** | Left junction PCR | F | oD33 | CGACAACATAAAACATCGAGTGC | 765 bp |
| | | R | oS314 | AGTGTGGGTCGAATGGGAAT | |
| | Right junction PCR | F | oS315 | AGTTCATCCGTCCAATCCCA | 413 bp |
| | | R | oD34 | TAATTCCAGCGAAAGGCAG | |
| **CeEcad-WF** | Left junction PCR | F | oS362 | CGCGAAATTCTACGGAAATC | 887 bp |
| **CeEcad-cpH** | | R | oS314 | AGTGTGGGTCGAATGGGAAT | |
| | Right junction PCR | F | oS315 | AGTTCATCCGTCCAATCCCA | 461 bp |
| | | R | oS363 | GGGAGAGAAAGGGAATTGTG | |
| | Three-primer PCR | F | oS360 | TCGAAATAACAGCCCTGATGG | 398 bp (WT); 561 bp (KI) |
| | | R | oS363 | GGGAGAGAAAGGGAATTGTG | |
| | | R | oS314 | AGTGTGGGTCGAATGGGAAT | |
| **CeEcad-C-WF** | Left junction PCR | F | oS360 | TCGAAATAACAGCCCTGATGG | 819 bp |
| | | R | oS314 | AGTGTGGGTCGAATGGGAAT | |
| | Right junction PCR | F | oS315 | AGTTCATCCGTCCAATCCCA | 595 bp |
| | | R | oD26 | AAGAGGGAAACGGGTAACAG | |
| **Vcl-WF** | Three-primer PCR | F | Vin-5F | CTTACATGGTGTCTACGTGG | 179 bp (KI) + 279 bp (WT) |
| **(mouse)** | | R | Vin-5R | TCGCGGTCAACAGGATTCAG | |
| | | R | Vin-3R | CTTGCTAGACCATTTCCGAG | |

**Table S3. Guide RNA and HDR donor sequences for *C. elegans* and mouse knock-in lines.**

| Strain | Item | Sequence / Parental strain |
|---|---|---|
| **CeEcad-WF** | crRNA (5′→3′) | CCTGTGTCGAGTGGCATTAC |
| | Parental strain | N2 |
| | HDR donor sequence (5′→3′)<br>Left homology arm –<br>Knock-in fragment –<br>Right homology arm | TCGAAATAACAGCCCTGATGGCGACTACAAACTACAAACTACA<br>AACTATAATAACCCCCAATTTTTTTCAGTACTCAATGGCCGGCC<br>TACGTAAACCAGTAGGAGGTTCCGGAGGTGTCCTCACCGAGG<br>TCGAGATGGACCACTACCGTGAGCCATTCCTCAACCCAGTCGA<br>CCGTGAGCCACTCTGGCGTTTCCCAAACGAGCTCCCAATCGC<br>CGGAGAGCCAGCCAACATCGTCGCCCTCGTCGAGGAGTACAT<br>GGACTGGCTCCACCAATCCCCAGTCCCAAAGCTCCTCTTCTGG<br>GGAACCCCAGGAGTCCTCATCCCACCAGCCGAGGCCGCCCGT<br>CTCGCCAAGTCCCTCCCAAACTGCAAGGCCGTCGACATCGGA<br>CCAGGACTCAACCTCCTCCAAGAGGACAACCCAGACCTCATTG<br>GATCTGAGATCGCCCGTTGGCTCTCCACCCTCGAGATCTCCG<br>GAGGAGGAACTGGTGGATCTGGTGGAACCGGAGGTTCTGGAG<br>GTACTGGTGGATCCGCCGAGATCGGAACCGGATTCCCATTCG<br>ACCCACACTACGTCGAGGTCCTCGGAGAGCGTATGCACTACG<br>TCGACGTCGGACCACGTGACGGAACCCCAGTCCTCTTCCTCC<br>ACGGAAACCCAACCTCCTCCTACGTCTGGCGTAACATCATCCC<br>ACACGTCGCCCCAACCCACCGTTGCATCGCTCCAGATCTTATT<br>GGAATGGGAAAGTCCGACAAGCCAGACCTCGGATACTTCTTC<br>GACGACCACGTCCGTTTCATGGACGCCTTCATCGAGGCCCTC<br>GGACTCGAGGAGGTCGTCCTCGTCATCCACGACTGGGGATCC<br>GCCCTCGGATTCCACTGGGCCAAGCGTAACCCAGAGCGTGTC<br>AAGGGAATCGCCTTCATGGAGTTCATCCGTCCAATCCCAACCT<br>GGGACGAGTGGCCAGAGTTCGCCCGTGAGACCTTCCAAGCCT<br>TCCGTACCACCGACGTCGGACGTAAGCTCATCATCGACCAAAA<br>CGTCTTCATCGAGTGGACCCTCCCAATGGGAGTCTGGCGTGG<br>AGGATCTGGAGGAATGCCACTCGACACAGGAATGGGACCAGC<br>AATCGGAGGACACCCACCACACTACCCACCACGTGGAATGGC<br>GCCACCAAAAGATGATCATGAGCTGAACTCGAAGATCAAGGAT<br>CTTGAGACTGATCAGAATGCGGCACCGTACGATGAACTTCGGA<br>TCTACGACGATGAGCGGGACAATA |
| **CeEcad-cpH** | crRNA (5′→3′) | GGACCCTCCCAATGGGAGTC |
| | Parental strain | CeEcad-WF |
| | HDR donor sequence (5′→3′)<br>Mutation edits | TCCTCCAGATCCTCCACGGACGACTCCCATTGGGAGGGTTCC<br>CTCGATGAAGACGTTTTGGTCGATGATGAGCTTACGTCC |
| **CeEcad-C-WF** | crRNA (5′→3′) | CGAAAGTGCCCAATAAACGA |
| | Parental strain | N2 |
| | HDR donor sequence (5′→3′)<br>Left homology arm –<br>Knock-in fragment –<br>Right homology arm | ATGGCCGGCCTACGTAAACCAGTAATGCCACTCGACACAGGA<br>ATGGGACCAGCAATCGGAGGACACCCACCACACTACCCACCA<br>CGTGGAATGGCGCCACCAAAAGATGATCATGAGCTGAACTCG<br>AAGATCAAGGATCTTGAGACTGATCAGAATGCGGCACCGTACG<br>ATGAACTTCGGATCTACGACGATGAGCGGGACAATATTTCTGT<br>CGTCACGTTGGAGAGTATCGAAAGTGCCCAAGGAGGCTCTGG<br>GGGTTCAGGCGGATCGGGTGGGTCCGGAGGCTCAGGGGGTT<br>CTGTCCTCACCGAGGTCGAGATGGACCACTACCGTGAGCCAT<br>TCCTCAACCCAGTCGACCGTGAGCCACTCTGGCGTTTCCCAAA<br>CGAGCTCCCAATCGCCGGAGAGCCAGCCAACATCGTCGCCCT<br>CGTCGAGGAGTACATGGACTGGCTCCACCAATCCCCAGTCCC<br>AAAGCTCCTCTTCTGGGGAACCCCAGGAGTCCTCATCCCACCA<br>GCCGAGGCCGCCCGTCTCGCCAAGTCCCTCCCAAACTGCAAG<br>GCCGTCGACATCGGACCAGGACTCAACCTCCTCCAAGAGGAC<br>AACCCAGACCTCATTGGATCTGAGATCGCCCGTTGGCTCTCCA<br>CCCTCGAGATCTCCGGAGGAGGAACTGGTGGATCTGGTGGAA |

| Strain | Item | Sequence / Parental strain |
|---|---|---|
| | | CCGGAGGTTCTGGAGGTACTGGTGGATCCGCCGAGATCGGAA<br>CCGGATTCCCATTCGACCCACACTACGTCGAGGTCCTCGGAG<br>AGCGTATGCACTACGTCGACGTCGGACCACGTGACGGAACCC<br>CAGTCCTCTTCCTCCACGGAAACCCAACCTCCTCCTACGTCTG<br>GCGTAACATCATCCCACACGTCGCCCCAACCCACCGTTGCATC<br>GCTCCAGATCTTATTGGAATGGGAAAGTCCGACAAGCCAGACC<br>TCGGATACTTCTTCGACGACCACGTCCGTTTCATGGACGCCTT<br>CATCGAGGCCCTCGGACTCGAGGAGGTCGTCCTCGTCATCCA<br>CGACTGGGGATCCGCCCTCGGATTCCACTGGGCCAAGCGTAA<br>CCCAGAGCGTGTCAAGGGAATCGCCTTCATGGAGTTCATCCG<br>TCCAATCCCAACCTGGGACGAGTGGCCAGAGTTCGCCCGTGA<br>GACCTTCCAAGCCTTCCGTACCACCGACGTCGGACGTAAGCT<br>CATCATCGACCAAAACGTCTTCATCGAGTGGACCCTCCCAATG<br>GGAGTCTGGCGTTAAACGACGGACTATATTATCATCTCTTTTTT<br>TCTGATCTTTTTTCGCGCTAATTTTTCACAATTCCCTTTCTCTCC<br>CCACCATCTCAATATTCCCTGCGATACTCGTCAACCGGTGTCA<br>AATCACACACACACAAAAATCTGCACTTTTTGACTCCTATCTGT<br>TACCTAGAACGGGTTCTGTTTTCTTGTTATAGGGCCCTTTTTAC<br>CCCAAAATTGTACACATTTTTTATGCAAGATTATGCAAGATTTTC<br>TAAC |
| **CeVin-WF** | crRNA (5′→3′) | GTGGTGAGGCATTGACACTT |
| | Parental strain | N2 |
| | HDR donor sequence (5′→3′)<br>Left homology arm –<br>Knock-in fragment –<br>Right homology arm | CCCTCCAAGATCGCCTCTCTCCCGGAAAAAATTAGAAATGATA<br>AGAAAATAAGACAAGGAACACAACTTTACAGCAATAGTTTAGTT<br>TTCAAGTTTTCCAAAAATCTACTAAAAAAAACTATTTTTCCAGCC<br>ACCACCATCATCCCAAGTGATCCGAGGAGGTTCCGGAGGTGT<br>CCTCACCGAGGTCGAGATGGACCACTACCGTGAGCCATTCCT<br>CAACCCAGTCGACCGTGAGCCACTCTGGCGTTTCCCAAACGA<br>GCTCCCAATCGCCGGAGAGCCAGCCAACATCGTCGCCCTCGT<br>CGAGGAGTACATGGACTGGCTCCACCAATCCCCAGTCCCAAA<br>GCTCCTCTTCTGGGGAACCCCAGGAGTCCTCATCCCACCAGC<br>CGAGGCCGCCCGTCTCGCCAAGTCCCTCCCAAACTGCAAGGC<br>CGTCGACATCGGACCAGGACTCAACCTCCTCCAAGAGGACAA<br>CCCAGACCTCATTGGATCTGAGATCGCCCGTTGGCTCTCCACC<br>CTCGAGATCTCCGGAGGAGGAACTGGTGGATCTGGTGGAACC<br>GGAGGTTCTGGAGGTACTGGTGGATCCGCCGAGATCGGAACC<br>GGATTCCCATTCGACCCACACTACGTCGAGGTCCTCGGAGAG<br>CGTATGCACTACGTCGACGTCGGACCACGTGACGGAACCCCA<br>GTCCTCTTCCTCCACGGAAACCCAACCTCCTCCTACGTCTGGC<br>GTAACATCATCCCACACGTCGCCCCAACCCACCGTTGCATCGC<br>TCCAGATCTTATTGGAATGGGAAAGTCCGACAAGCCAGACCTC<br>GGATACTTCTTCGACGACCACGTCCGTTTCATGGACGCCTTCA<br>TCGAGGCCCTCGGACTCGAGGAGGTCGTCCTCGTCATCCACG<br>ACTGGGGATCCGCCCTCGGATTCCACTGGGCCAAGCGTAACC<br>CAGAGCGTGTCAAGGGAATCGCCTTCATGGAGTTCATCCGTC<br>CAATCCCAACCTGGGACGAGTGGCCAGAGTTCGCCCGTGAGA<br>CCTTCCAAGCCTTCCGTACCACCGACGTCGGACGTAAGCTCAT<br>CATCGACCAAAACGTCTTCATCGAGTGGACCCTCCCAATGGGA<br>GTCTGGCGTGGAGGATCTGGAGGAAGTGTCAATGCCTCACCA<br>CCAACAGCTCCAATCATTCACAATAAGATGATCATCCGAGAAG<br>ATATTCCAGCTCCACCACG |
| **CeVin-cpH** | crRNA (5′→3′) | GGACCCTCCCAATGGGAGTC |
| | Parental strain | CeVin-WF |
| | HDR donor sequence (5′→3′)<br>Mutation edits | TCCTCCAGATCCTCCACGGACGACTCCCATTGGGAGGGTTCC<br>CTCGATGAAGACGTTTTGGTCGATGATGAGCTTACGTCC |

| Strain | Item | Sequence / Parental strain |
|---|---|---|
| **CeVin-C-WF** | crRNA (5′→3′) | ATGAACGGGGATTTAGAAGT |
| | Parental strain | N2 |
| | HDR donor sequence (5′→3′)<br>Left homology arm – Knock-in fragment – Right homology arm | GAATTTCAGGCAGCGAAGAAGATGATGAAGCAATGCAACAATT<br>GGTGCATAATGCTCAAAACTTGATGCAATCTGTGAAGGATGTT<br>GTCCGTGCTGCTGAAGCCGCCTCTATTAAAATCCGTACCAACT<br>CGGGACTTCGTCTCCGTTGGCTCCGAAAGCCAATGTGGTCCA<br>ACTTCGGAGGCTCTGGGGGTTCAGGCGGATCGGGTGGGTCC<br>GGAGGCTCAGGGGGTTCTGTCCTCACCGAGGTCGAGATGGAC<br>CACTACCGTGAGCCATTCCTCAACCCAGTCGACCGTGAGCCA<br>CTCTGGCGTTTCCCAAACGAGCTCCCAATCGCCGGAGAGCCA<br>GCCAACATCGTCGCCCTCGTCGAGGAGTACATGGACTGGCTC<br>CACCAATCCCCAGTCCCAAAGCTCCTCTTCTGGGGAACCCCA<br>GGAGTCCTCATCCCACCAGCCGAGGCCGCCCGTCTCGCCAAG<br>TCCCTCCCAAACTGCAAGGCCGTCGACATCGGACCAGGACTC<br>AACCTCCTCCAAGAGGACAACCCAGACCTCATTGGATCTGAGA<br>TCGCCCGTTGGCTCTCCACCCTCGAGATCTCCGGAGGAGGAA<br>CTGGTGGATCTGGTGGAACCGGAGGTTCTGGAGGTACTGGTG<br>GATCCGCCGAGATCGGAACCGGATTCCCATTCGACCCACACT<br>ACGTCGAGGTCCTCGGAGAGCGTATGCACTACGTCGACGTCG<br>GACCACGTGACGGAACCCCAGTCCTCTTCCTCCACGGAAACC<br>CAACCTCCTCCTACGTCTGGCGTAACATCATCCCACACGTCGC<br>CCCAACCCACCGTTGCATCGCTCCAGATCTTATTGGAATGGGA<br>AAGTCCGACAAGCCAGACCTCGGATACTTCTTCGACGACCAC<br>GTCCGTTTCATGGACGCCTTCATCGAGGCCCTCGGACTCGAG<br>GAGGTCGTCCTCGTCATCCACGACTGGGGATCCGCCCTCGGA<br>TTCCACTGGGCCAAGCGTAACCCAGAGCGTGTCAAGGGAATC<br>GCCTTCATGGAGTTCATCCGTCCAATCCCAACCTGGGACGAGT<br>GGCCAGAGTTCGCCCGTGAGACCTTCCAAGCCTTCCGTACCA<br>CCGACGTCGGACGTAAGCTCATCATCGACCAAAACGTCTTCAT<br>CGAGTGGACCCTCCCAATGGGAGTCTGGCGTTAAATCCCCGT<br>TCATTACCGATTCTCTTTCTTCTCAATCAATGTTTTGCATATTTT<br>ATGTATTTTTCTACTGCTTTGCTAGTCATTTAAATTTTTTGCATA<br>ATAATTTTTTTTCATTCTTAATAATGTTACTTTTCAGTTTGAAATT<br>GCTCCAAATCTTCCACTCTCCTTTCCCCATG |
| **CeLam-WF** | crRNA (5′→3′) | AATGTGTAACAGTCATCGCA |
| | Parental strain | N2 |
| | HDR donor sequence (5′→3′)<br>Left homology arm – Knock-in fragment – Right homology arm | TGTCAACCATGCGATTGTGAATACATCGGATCCGAGAACCAAC<br>AATGTGATGTCAATTCTGGACAATGCTTGTGTAAGGAGAATGTT<br>GAAGGAAGAAGATGCGACCAGTGCGCTGAAAACCGATATGGA<br>ATCACTCAAGGATGCTTGCCATGCGGAGGTTCCGGAGGTGTC<br>CTCACCGAGGTCGAGATGGACCACTACCGTGAGCCATTCCTC<br>AACCCAGTCGACCGTGAGCCACTCTGGCGTTTCCCAAACGAG<br>CTCCCAATCGCCGGAGAGCCAGCCAACATCGTCGCCCTCGTC<br>GAGGAGTACATGGACTGGCTCCACCAATCCCCAGTCCCAAAG<br>CTCCTCTTCTGGGGAACCCCAGGAGTCCTCATCCCACCAGCC<br>GAGGCCGCCCGTCTCGCCAAGTCCCTCCCAAACTGCAAGGCC<br>GTCGACATCGGACCAGGACTCAACCTCCTCCAAGAGGACAAC<br>CCAGACCTCATTGGATCTGAGATCGCCCGTTGGCTCTCCACCC<br>TCGAGATCTCCGGAGGAGGAACTGGTGGATCTGGTGGAACCG<br>GAGGTTCTGGAGGTACTGGTGGATCCGCCGAGATCGGAACCG<br>GATTCCCATTCGACCCACACTACGTCGAGGTCCTCGGAGAGC<br>GTATGCACTACGTCGACGTCGGACCACGTGACGGAACCCCAG<br>TCCTCTTCCTCCACGGAAACCCAACCTCCTCCTACGTCTGGCG<br>TAACATCATCCCACACGTCGCCCCAACCCACCGTTGCATCGCT<br>CCAGATCTTATTGGAATGGGAAAGTCCGACAAGCCAGACCTCG<br>GATACTTCTTCGACGACCACGTCCGTTTCATGGACGCCTTCAT<br>CGAGGCCCTCGGACTCGAGGAGGTCGTCCTCGTCATCCACGA<br>CTGGGGATCCGCCCTCGGATTCCACTGGGCCAAGCGTAACCC |

| Strain | Item | Sequence / Parental strain |
|---|---|---|
| | | AGAGCGTGTCAAGGGAATCGCCTTCATGGAGTTCATCCGTCCA<br>ATCCCAACCTGGGACGAGTGGCCAGAGTTCGCCCGTGAGACC<br>TTCCAAGCCTTCCGTACCACCGACGTCGGACGTAAGCTCATCA<br>TCGACCAAAACGTCTTCATCGAGTGGACCCTCCCAATGGGAGT<br>CTGGCGTGGAGGATCTGGAGGAGATGACTGTTACACATTGAT<br>CCAGAGCCGTGTTAATGTGTTCAGAGAGAAAGTTAAAAGTCTT<br>GATAATACTCTTCAAGAGATTATTGAAAACCCAGCTCCAGTGAA<br>TGACACCAAATTTGATGAGAAAGTCAAGGAAACGTCTCGTGCT<br>GCTTCAGAAG |
| **CeLam-cpH** | crRNA (5′→3′) | GGACCCTCCCAATGGGAGTC |
| | Parental strain | CeLam-WF |
| | HDR donor sequence (5′→3′)<br>Mutation edits | TCCTCCAGATCCTCCACGGACGACTCCCATTGGGAGGGTTCC<br>CTCGATGAAGACGTTTTGGTCGATGATGAGCTTACGTCC |
| **CeLam-C-WF** | crRNA (5′→3′) | GTCATCAATTTGGAGCAAGA |
| | Parental strain | N2 |
| | HDR donor sequence (5′→3′)<br><br>Left homology arm –<br>Knock-in fragment –<br>Right homology arm<br><br>Mutations to avoid sgRNA targeting | TCAGCAATACCGCGCTGACGAGGACGTAAAGGTGGCACAGCT<br>CAAAAATGATATTTCTGAGCTCCAGAAGGAGGTTTGTTTTGCTG<br>GCTATAGTATTCGTATAATTAAACTGTATTTTTCAGGTTTTGAAC<br>CTAGAGGAAATCCGTGACAACTTGCCAACCAAATGCTTCAATG<br>TCATCAATTTGGAACAGGAAGGACAGAAAGGAGCTTCCGGCG<br>CCTCAGGGGCAAGCGGTGCGTCTGGAGCTAGTGGCGCCTCG<br>GTCCTCACCGAGGTCGAGATGGACCACTACCGTGAGCCATTC<br>CTCAACCCAGTCGACCGTGAGCCACTCTGGCGTTTCCCAAAC<br>GAGCTCCCAATCGCCGGAGAGCCAGCCAACATCGTCGCCCTC<br>GTCGAGGAGTACATGGACTGGCTCCACCAATCCCCAGTCCCA<br>AAGCTCCTCTTCTGGGGAACCCCAGGAGTCCTCATCCCACCA<br>GCCGAGGCCGCCCGTCTCGCCAAGTCCCTCCCAAACTGCAAG<br>GCCGTCGACATCGGACCAGGACTCAACCTCCTCCAAGAGGAC<br>AACCCAGACCTCATTGGATCTGAGATCGCCCGTTGGCTCTCCA<br>CCCTCGAGATCTCCGGAGGAGGAACTGGTGGATCTGGTGGAA<br>CCGGAGGTTCTGGAGGTACTGGTGGATCCGCCGAGATCGGAA<br>CCGGATTCCCATTCGACCCACACTACGTCGAGGTCCTCGGAG<br>AGCGTATGCACTACGTCGACGTCGGACCACGTGACGGAACCC<br>CAGTCCTCTTCCTCCACGGAAACCCAACCTCCTCCTACGTCTG<br>GCGTAACATCATCCCACACGTCGCCCCAACCCACCGTTGCATC<br>GCTCCAGATCTTATTGGAATGGGAAAGTCCGACAAGCCAGACC<br>TCGGATACTTCTTCGACGACCACGTCCGTTTCATGGACGCCTT<br>CATCGAGGCCCTCGGACTCGAGGAGGTCGTCCTCGTCATCCA<br>CGACTGGGGATCCGCCCTCGGATTCCACTGGGCCAAGCGTAA<br>CCCAGAGCGTGTCAAGGGAATCGCCTTCATGGAGTTCATCCG<br>TCCAATCCCAACCTGGGACGAGTGGCCAGAGTTCGCCCGTGA<br>GACCTTCCAAGCCTTCCGTACCACCGACGTCGGACGTAAGCT<br>CATCATCGACCAAAACGTCTTCATCGAGTGGACCCTCCCAATG<br>GGAGTCTGGCGTTAAGATAATCTTTATAATTTTTCTATCAAACTT<br>TTTCTATGAAAATCGATCAGATAATTTATTAACTCTAATATTAAA<br>ATATCTGAAAAAGAGCCCTTAAAAATTATTAGGTCGGCAAATAA<br>TTATTGACGTGTTTTGCCAATTAACGATTTTTTGTAAATTTTATA<br>TGTGTCTCTTACAGTAAACCGAGAGTGTATAAAAGCATATGAAT<br>ATCAGTTGCTTTGGTCATCTGAGTAGTAAT |
| **Vcl-WF (mouse)** | sgRNA (5′→3′) | TTTGTAGCTAACTGATGAGC |
| | Parental strain | FVB/N |
| | HDR donor sequence (5′→3′)<br><br>Left homology arm – | TTGGGTGCTCAGGATCAACTAGGGTCTTCATGCTGGCATGGCA<br>AGTACTCTCCTGTCTCCTTAGACCAAGAAATGGTTCTGTGTTTT<br>TAAATCTGATTCGAATTTCAAATTTTTAAAACTTAAAAAAAAATT<br>TATGCGTATGGATGCGTCGCACTGCTCGTATGCCTAGTGCCCT |

| Strain | Item | Sequence / Parental strain |
|---|---|---|
| | Knock-in fragment – Right homology arm | CAGAACCCAGAAGAGGGTATTGGATGCCCTGGAACCGCAGTT<br>GTAGATGGTTGTGAGCAGCCATGTGGATGCTGGGAATTTGAAC<br>CCATAGTTTCTGGCGGAGCAACAAGTGCTCTTAACTGCTCAGC<br>CATCTCTCTAGGCCCTCAAATCTTTTCGTAATTCTTTCCCATCA<br>AGTAGCATGAATACATTTATCATGTATCATCTTTCCCATGATGT<br>ACTTGGGGAGTATCTTTTAGTTATAGATCCTAGAAACAAAAACA<br>TTGTGTAATGATCATGTACACAAGTATTTACTCCAAGTAATAAG<br>TCTAGAAACACGTTAAGACAAGAATTATATTTACTTCCTTTGGT<br>ATGTTTTCTTATTTAATGTCAGTAAAAGAAGTCAGGTTGGTTAA<br>TGACTAAATTGGGTCACTATACATAAGTAGGATTGGTGGTTCC<br>AGTTTTCAACATTTCTTTACACTTTCCAATGAGGCTTTTGATGAT<br>GAGATTCCTGGTAAGTGGCTTCTTTGAGTCATGACATTTAGAAT<br>TAGGGACATTGAGATCACATGGCAGATGTAGAACTTGAGGCTC<br>CAGTGGGAAGTTACTGACTCTCATTTGCCCTTTGACAACCAGA<br>ACCAGAACCAGGTCTTAAATCTTCTATGGGGATTGGCTTTGGG<br>AATCGCTGATCTGGTAATGTTATGTATGCCTCCTATGGGATATG<br>AGCAACACATAGCTGCTGGTTTCTTATAGAGAAAAAAGTTTTTA<br>TTGATATTGCACAGATGTCCTCATGGTTCAGTGTTGATGGTCTT<br>ACATGGTGTTTTTATTGATATTGCACAGATGTCGTCATGGTTCA<br>GTGTTGGTGGTCTTACATGGTGTCTACGTGGGCCCATTTCCTA<br>GGCTGGTTTTGGTAGTAGCTTCCTGGGCTTTACTGACAACCTG<br>CTTTATTCACATTTGTAGCTAACTGATGGAGGTTCTGGAGGAG<br>TGCTGACTGAAGTCGAGATGGACCATTACCGCGAGCCGTTCC<br>TGAATCCTGTTGACCGCGAGCCACTGTGGCGCTTCCCAAACG<br>AGCTGCCAATCGCCGGTGAGCCAGCGAACATCGTCGCGCTGG<br>TCGAAGAATACATGGACTGGCTGCACCAGTCCCCTGTCCCGA<br>AGCTGCTGTTCTGGGGCACCCCAGGCGTTCTGATCCCACCGG<br>CCGAAGCCGCTCGCCTGGCCAAAAGCCTGCCTAACTGCAAGG<br>CTGTGGACATCGGCCCGGGTCTGAATCTGCTGCAAGAAGACA<br>ACCCGGACCTGATCGGCAGCGAGATCGCGCGCTGGCTGTCG<br>ACGCTCGAGATTTCCGGCGGAGGAACAGGTGGTTCTGGTGGA<br>ACAGGGGGTAGCGGAGGTACAGGAGGAAGTGCAGAAATCGG<br>TACTGGCTTTCCATTCGACCCCCATTATGTGGAAGTCCTGGGC<br>GAGCGCATGCACTACGTCGATGTTGGTCCGCGCGATGGCACC<br>CCTGTGCTGTTCCTGCACGGTAACCCGACCTCCTCCTACGTGT<br>GGCGCAACATCATCCCGCATGTTGCTCCGACCCATCGCTGCA<br>TTGCTCCAGACCTGATCGGTATGGGCAAATCCGACAAACCAGA<br>CCTGGGTTATTTCTTCGACGACCACGTCCGCTTCATGGATGCC<br>TTCATCGAAGCCCTGGGTCTGGAAGAGGTCGTCCTGGTCATTC<br>ACGACTGGGGCTCCGCTCTGGGTTTCCACTGGGCCAAGCGCA<br>ATCCAGAGCGCGTCAAAGGTATTGCATTTATGGAGTTCATCCG<br>CCCTATCCCGACCTGGGACGAATGGCCAGAATTTGCCCGCGA<br>GACCTTCCAGGCCTTCCGCACCACCGACGTCGGCCGCAAGCT<br>GATCATCGATCAGAACGTTTTTATCGAGTGGACGCTGCCGATG<br>GGTGTCTGGCGCGGTGGATCTGGTGGAGAGCTGGCTCCTCCT<br>AAGCCACCTCTGCCTGAGGGTGAAGTCCCTCCACCCAGGCCC<br>CCACCACCAGAAGAGAAGGATGAAGAGTTCCCTGAGCAGAAA<br>GCTGGTGAGGTGATTAACCAGCCAATGATGATGGCCGCCAGG<br>CAGCTCCACGATGAAGCTCGGAAATGGTCTAGCAAGGTAAGC<br>CCCGGGACTTTCTTTCCTTGCTGGGGATGCGTGCCTTTTCAGA<br>AAGGACGGAGAACACCCCAGGAAGAAGTCCATGTGATCACTTT<br>GCTTCTTGCTCAGTTCCTTCATTTCAGGCAGTGGGTTTTTACCA<br>GCGATTTAGTGAGAAGTCTCTAGTGTTTGATGGCTTGCCTTTG<br>CCTGGGCAGAGGTAAAAGCTGAGATGTCAGTGGCTAAAGAAG<br>AGAGTCTGGCCCCTTTGTCCAGTAGAGTGTCAGCTGTACTGAT<br>CCTGACATCTGGGGTCTTTTTTAGAAACTGTTTATCCTTTCTCA<br>GAATCATCTTGTGCTTTTTCTCTACTTAGGAATGCTCCAGGTTA<br>TGTTCAGCTCTGATTATTTAAAATTCTAACTATTCCTGTGGCTG<br>AGATCTGGGTGCAATCCAGTTCTACTGCATGCTCACCTTTTCC<br>CATCACTGGAAATGAGTAGTTGGTTTTTTTTGCTTAGAGCAAAA |

| Strain | Item | Sequence / Parental strain |
|---|---|---|
| | | TCCTGCTGCTCATCTGCCAATGAGGATATGGTTCTTAGCTCGA<br>GACTGTAAGGAATCAGAAATCTAGCTCTTCATTCCCACTATATT<br>CCTTTATATTCATCTTAACAGAATATACTTCCCAGAGGAAGTTA<br>ATCTTGTGTTGTTGTTGGATTTTTTCTTCAGGCTGAGAAATTGA<br>CCTTGGTTTGTTCTATTTTGTTTGTTTCTTGTTTTGTTTTTCAAG<br>ACATGGTTTTTCTGTGTAGCTCTGGCTACCCTAGAACTCAACCT<br>GTACACCAGGCTGGCCTGGAACTCAAGAGATCCCGCCTGCCT<br>CTTTTCCTGAGTGCTGAGATTAAAGGCAAGTGCCACCACTGCC<br>AATGTTACCTTGGAGCCCAGTCTGGTGTTGGATCTGACCAGTG<br>GAATTATTTATAATGGTGTCTCTCCAACACTCCCAGGATCATGT<br>AGCATAATCCTACTGTGAATGGACAATGGGTGACAATAGGGTT<br>GATTTCCCCCAAAGAAAATGCAAATGTAGTTATAATGAGCCTG<br>CACATATTGTGAAAATGTATGTGGAATATAAAGGGTCAGACAG<br>CACTGGCCCATATAGTTGC |
